# Environmental niche and demographic modeling of American chestnut near its southwestern range limit

**DOI:** 10.1101/2021.09.10.459451

**Authors:** Robert G. Laport, Zoe S. Brookover, Brian D. Christman, Julienne Ng, Kevin Philley, J. Hill Craddock

**Affiliations:** Department of Biology, Rhodes College, Memphis 38112; United States Army Corps of Engineers, Engineer Research and Development Center, Vicksburg 39180; Department of Biology, Geology, and Environmental Science, University of Tennessee at Chattanooga, Chattanooga 37403

**Keywords:** *Castanea dentata*, population matrix model, population size structure, restoration, species distribution modeling

## Abstract

The inadvertent introduction and rapid spread of chestnut blight (caused by *Cryphonectria parasitica* (Murr.) Barr) in the early 20th century resulted in the demise of American chestnut (*Castanea dentata* (Marsh.) Borkh.; Fagaceae) as a major component of forest canopies and had negative impacts on eastern forest communities. Research efforts over the last century have documented the persistence of occasional trees and root crown/stump sprouts throughout much of the species’ historic range providing the basis for ongoing breeding of blight-resistant stock and restoration efforts. Unfortunately, it remains unclear how much of the historic range remains climatically suitable for remnant trees that may harbor unique genetic variation for successful reintroduction efforts. Here we investigate whether the southwestern portion of the historical range remains environmentally suitable for undiscovered remnant populations of *C. dentata* using environmental niche modeling. We also use stage-structured matrix projection models to investigate the potential demographic future of *C. dentata* in W Tennessee, N Mississippi, SW Kentucky, and NW Alabama based upon observations of American chestnut in these areas over the last several decades. We found that suitable habitat associated with higher elevations and areas of high forest canopy cover occurs throughout much of the southwestern portion of the historical range and that populations of American chestnut in these areas are predicted to drastically decline over the next ~100-200 years without conservation interventions to mitigate the negative consequences of chestnut blight.

## Introduction

The unintentional introduction and rapid spread of chestnut blight (*Cryphonectria parasitica* (Murr.) Barr) in the early 20^th^ century decimated American chestnut (*Castanea dentata* (Marsh.) Borkh.; Fagaceae) populations (Ziegler, 1920; Smith, 2000; Jacobs *et al.*, 2013; Dalgleish *et al.*, 2016). The loss of this species as a dominant canopy tree has profoundly impacted forest communities and the ecological dynamics of eastern hardwood forests. For example, the decline of annual large, edible seed production has impacted plant-animal interactions and wildlife population dynamics (Diamond *et al.*, 2000; Dalgleish and Swihart, 2012), and the loss of a major canopy tree has altered forest community dynamics, nutrient cycling, and fire regimes (Brooks, 1937, McCormick and Platt, 1980, Ellison *et al.*, 2005; Elliot and Swank, 2008, Jacobs *et al.*, 2013; De Bruijn *et al.*, 2014; Gustafson *et al.*, 2017). Given its previously widespread distribution, rapid demise, and ecological significance to the forests of eastern North America, policy and research efforts have been focused on the ultimate goal of breeding blight resistant trees and restoring American chestnut to its former range.

Research efforts since the middle of the 20^th^ century aimed at conserving *C. dentata* have documented the persistence of root crown/stump sprouts, saplings, and occasional fruiting trees throughout much of the American chestnut’s former distribution. For example, remnant populations of trees have been documented in parts of Connecticut (Stephens and Waggoner, 1980; Paillet, 1982, 2002), Massachusetts (Paillet, 1988, 2002), Virginia (Stephenson *et al.*, 1991), Ohio (Schwadron, 1995), southern Ontario (Tindall *et al.*, 2004, Van Drunen *et al.*, 2017), western New York (Laport, 2020), and southwest Tennessee (Laport *et al.*, 2020). Remnant trees are also continually being documented on crowd-sourcing mobile applications such as iNaturalist (www.inaturalist.org) and TreeSnap (www.treesnap.org), with many of these observations being archived by the Global Biodiversity Information Facility (GBIF; www.gbif.org). These natural history observations suggest the highest densities of remnant trees occur throughout the Cumberland Plateau, Appalachian Mountains, and the northeastern US. Fewer contemporary *C. dentata* occurrences are known from the southwestern range margin compared to core distribution populations, but it is not clear to what degree this disparity reflects actual differences in available habitat, population occurrence density, or whether this is because some parts of the range have been under-surveyed (Jacobs *et al.*, 2013; Dalgleish *et al.*, 2016).

Ecological/Environmental Niche Modeling (ENM) or Species Distribution Modeling (SDM) approaches provide one approach to identify areas that are environmentally suitable for remnant American chestnut, and to aid conservation and reintroduction efforts. Recent studies have implemented ENM/SDM approaches to investigate the current and future abiotic niche of remnant American chestnut with the goal of identifying areas of ecological suitability throughout the historical range to inform future restoration efforts. For example, models constructed using known occurrences of American chestnut in Shenandoah National Park and Mammoth Cave National Park projected suitable habitat in areas of relatively undisturbed upland forest, often on slopes or at relatively high elevations. These studies also found that young forests on previously cultivated lands were almost universally predicted as unsuitable for chestnut occurrence (Fei *et al.*, 2007; Santoro, 2013). Another recent study using ENM/SDM approaches examined the projected abiotic habitat suitability for American chestnut under multiple scenarios of climate change over the next several decades. At a continental scale, this study found that suitable habitat would shift northward into the northeastern US and southern Canada (Barnes and Delborne, 2019). These studies provide critical information for guiding the discovery, protection, and reintroduction of American chestnut in mountainous core parts of its historical range where land use history resulted in the preservation of relatively large areas of suitable habitat, or at the northern range margin where the species may be expanding into new habitat in response to climate change. Unfortunately, these previous models offer only a limited window into the ecology of the species near the southern and western historical range limits by relying principally upon occurrences from core parts of the range or by investigating continental scale patterns that paint in broad strokes.

Populations occurring near species’ range limits may inhabit marginal ecological conditions not found within core parts of the distribution, which could promote the evolution and maintenance of novel genetic variation and local adaptation (Halbritter *et al.*, 2015; Angert *et al.*, 2020). In addition to representing populations that may be locally adapted to the unique environments of the Gulf Coastal Plain physiographic region, populations of *C. dentata* near the southern and western historical range limits are also hypothesized to potentially harbor genetic diversity more closely reflecting the ancestral populations that expanded from refugia following the end of the Wisconsin glacial maximum (Russell, 1987; Huang, 1998; Kubisiak and Roberds, 2006, Dane, 2009). This genetic diversity could greatly aid breeding and restoration efforts under a rapidly changing climate that is expected to push suitable habitat farther northward (Huang *et al.*, 1998; Shaw *et al.*, 2012; Barnes and Delborne, 2019), but could also shed light on the evolution and ecology of a species that was nearly extirpated prior to extensive study with modern ecological methods.

In addition to experiencing atypical or marginal environments, populations near species range limits also experience unique demographic processes arising, in part, from the challenges of small population sizes (*e.g.*, mate limitation and inbreeding depression, local adaptation and outbreeding depression, mutational load; Ellstrand and Elam, 1993; Angert *et al.*, 2020; Perrier *et al.*, 2020). Range-edge populations can become locally adapted via the emergence of novel genetic variation, but can also be demographically unstable, or transient, limiting their demographic footprint (*e.g.*, Savolainen *et al.*, 2007; Angert *et al.*, 2008, 2020). Remnant populations of *C. dentata* occurring in the southwestern portion of the historical range have previously been noted to have higher levels of genetic variation than populations in other parts of the distribution, potentially reflecting local adaptation or that these populations may more closely reflect the genetic variation from populations that expanded from glacial refugia following the last glacial maximum (Russell, 1987; Huang *et al.*, 1998, Shaw *et al.*, 2012). Prior evidence also suggests that populations in the southwestern part of the historical distribution are relatively small in comparison to the rest of the distribution and may have been afflicted by *Phytophthora cinnamomi* (Rands) prior to the advent of chestnut blight (Anagnostakis, 2001; Dalgleish *et al.*, 2016). Yet, relatively little is known about the population dynamics of the species from this region and few studies have investigated American chestnut population dynamics near its historical range margins. The demography of American chestnut has been studied near its northern distributional limit in southern Ontario (Van Drunen, 2018) and central Maine (Elwood, 2014), as well as beyond the historical range in naturalized populations in Michigan and Wisconsin (Davelos and Jarosz, 2004; Stevens *et al.*, 2014). These studies reveal that in the absence of chestnut blight, or when blight affliction is modest, populations of chestnut may remain stable or expand into new habitat. It is unclear how the population dynamics of *C. dentata* near its southwestern range margin compare to the demography of other remnant populations near or beyond the native range, and whether such populations (and their novel genetic variation) are at increased risk of local extinction. Recently, small populations near and beyond the historical southwestern range limit in SW Tennessee and adjacent N Mississippi have been documented (Laport *et al.*, 2020) suggesting the demography of the species in this region remains poorly understood, particularly with respect to the prevalence of blight and population persistence.

The paucity of ecological and demographic information for American chestnut populations throughout W Tennessee, N Mississippi, SW Kentucky, and NW Alabama makes it difficult to extend current research, breeding, and reintroduction efforts to southwestern parts of the historical range. Yet, such populations are historically important components of the native forest communities, and may also be extremely valuable sources of genetic variation for ongoing restoration efforts that could increase the potential for successful regional reintroductions or for mitigating the effects of climate change on reintroduced populations. Here we make an initial attempt to (1) identify whether environmentally suitable habitat currently exists near the historical southwestern range margin that could support undiscovered remnant populations of *C. dentata* using ecological/environmental niche modeling approaches, and (2) to model the demography of *C. dentata* in W Tennessee, N Mississippi, SW Kentucky, and NW Alabama using simple projection matrix models to better understand the current status and future outlook for persistence of American chestnut throughout the southwestern portion of its range. We found that suitable habitat occurs throughout much of the southwestern range extent, but is most strongly predicted to coincide with higher elevations and areas of high forest canopy cover. Additionally, remnant populations of American chestnut in W Tennessee, N Mississippi, SW Kentucky, and NW Alabama are predicted to drastically decline in number over the next ~100–200 years, representing a significant loss to the local forest communities, as well as a loss of potentially locally adapted genomes that may be crucial for breeding and restoration efforts without conservation interventions.

## Methods

### ENVIRONMENTAL NICHE MODELS

#### Population occurrence data

We obtained occurrence information for American chestnut throughout W Tennessee, N Mississippi, SW Kentucky, and NW Alabama from citizen science reports (iNaturalist), GBIF records (https://doi.org/10.15468/dl.gqv45a), and author-identified occurrences. Additional Mississippi records were obtained from herbarium records at the University of Mississippi (MISS), University of Southern Mississippi (USMS), Mississippi State University (MISSA), Northern Kentucky University (KNK), University of North Carolina at Chapel Hill (NCU), Louisiana State University (LSU), and the Mississippi Museum of Natural Science (MMNS). We additionally obtained occurrence records throughout the study region from a database comprising >3,000 individual occurrences documented since the late 1990s and early 2000s by J. Schibig and collaborators (Schibig *et al.*, 2005, 2006). After filtering to an area encompassing W Tennessee, N Mississippi, SW Kentucky, NW Alabama, and NE Arkansas (NW corner: 37.3583°, −91.1444°, SE corner: 32.8416°, −85.6044°) we obtained 583 occurrences. We thinned the dataset to 140 occurrences by retaining only a single point in any given ~1km^2^ raster pixel (*i.e.*, removing all but a single point from within the same ~1 km^2^ raster pixel; see below) to reduce autocorrelation among clustered occurrences within populations (Table 1).

**Table 1.** --American chestnut observations and locality information throughout W. Tennessee, N. Mississippi, S.W. Kentucky, and N.W. Alabama used for environmental niche modeling. Observations included historical collections from herbaria, personal observations by the authors and colleagues, and engaged citizen-scientist observations from GBIF recorded and verified with the iNaturalist application. Detailed latitude/longitude location information is available upon request

| Date | Latitude | Longitude | County | State | Collectors | Collector Code/Voucher Accession |
| --- | --- | --- | --- | --- | --- | --- |
| 16-Jul-1969 | 34.76 | -88.43 | Alcorn | Mississippi | T. M. Pullen | MISS0018279 |
| 12-May-2019 | 35.03 | -88.73 | McNairy | Tennessee | R.G. Laport, D. Smith & J. Ng | SWMT05622 |
| 12-May-2019 | 35.03 | -88.74 | McNairy | Tennessee | R.G. Laport, D. Smith & J. Ng | SWMT05625 |
| 31-Oct-1981 | 33.91 | -89.26 | Calhoun | Mississippi | R. A. Stewart | MISS0018280 |
| 12-May-1967 | 34.11 | -89.50 | Calhoun | Mississippi | T. M. Pullen | MISS0018282 |
| 22-Jun-1980 | 33.29 | -89.10 | Choctaw | Mississippi | S. McDaniel | MMNS021400 |
| 15-Jun-1980 | 33.36 | -89.14 | Choctaw | Mississippi | S. McDaniel | MMNS021294 |
| 14-Jun-2013 | 33.42 | -89.26 | Choctaw | Mississippi | K. Philley | MMNS034856 |
| 21-Apr-1967 | 33.77 | -88.89 | Clay | Mississippi | T. R. Simmons, Jr | MISS0018284 |
| 22-Aug-1979 | 33.79 | -88.91 | Clay | Mississippi | K. E. Rogers | MMNS021663 |
| 14-Sep-2019 | 35.39 | -88.77 | Chester | Tennessee | R.G. Laport, D. Smith & J. Ng | SWMT05640 |
| 10-Nov-2019 | 35.38 | -88.77 | Chester | Tennessee | R.G. Laport, D. Smith & J. Ng | CSP19 |
| 1-May-1884 | 34.17 | -86.83 | Cullman | Alabama | C. Mohr | UNA00032071 |
| 6-Jun-2002 | 36.06 | -86.79 | Davidson | Tennessee | S. J. Baskauf | http://bioimages.vanderbilt.edu/ind-baskauf/11845#2002-06-06 |
| 15-Sep-1994 | 33.69 | -89.67 | Grenada | Mississippi | J. R. MacDonald | MISS0018287 |
| 20-Jun-1986 | 33.70 | -89.53 | Grenada | Mississippi | M. W. Morris | MISS_Morris2115 |
| 24-Apr-1994 | 33.71 | -89.69 | Grenada | Mississippi | J. R. MacDonald | MMNS013157 |
| 12-Jul-1980 | 33.73 | -89.76 | Grenada | Mississippi | R. A. Stewart II | MMNS004275 |
| 17-Jun-1986 | 33.75 | -89.58 | Grenada | Mississippi | M. W. Morris | MISS_Morris2107 |
| 27-Apr-1980 | 33.76 | -89.83 | Grenada | Mississippi | R. A. Stewart II | MMNS004504 |
| unknown | 36.31 | -88.26 | Henry | Tennessee | N/A | USDA_NPGS_3395807 |
| 12-Jun-2020 | 34.94 | -89.04 | Benton | Mississippi | R.G. Laport, D. Smith & J. Ng | SWMT05659 |
| 29-Aug-2004 | 36.75 | -86.23 | Allen | Kentucky | J. Schibig & M. Vance | KALL1 |
| 23-May-2004 | 36.66 | -87.35 | Christian | Kentucky | Caudell & J. Schibig | KCHR1 |
| 15-Jun-2004 | 36.76 | -88.43 | Marshall | Kentucky | J. Schibig & M. Vance | KMAR1 |
| 28-Jun-2003 | 36.73 | -85.87 | Monroe | Kentucky | J. Schibig & J. H. Craddock | KMON4 |
| 15-May-2003 | 36.72 | -88.05 | Trigg | Kentucky | J. H. Craddock | KTRI123 |
| 13-Aug-2003 | 36.71 | -88.07 | Trigg | Kentucky | J. Schibig & J. H. Craddock | KTRI148 |
| 26-May-2002 | 36.77 | -88.11 | Trigg | Kentucky | J. Schibig, Barber, J. H. Craddock & Torkelson | KTRI16 |
| 14-Jul-2002 | 36.72 | -88.08 | Trigg | Kentucky | J. Schibig | KTRI60 |
| 29-Sep-2002 | 36.78 | -88.11 | Trigg | Kentucky | J. Schibig, Barber, J. H. Craddock & Torkelson | KTRI86 |
| 3-May-2003 | 36.76 | -88.11 | Trigg | Kentucky | J. Schibig & M. Vance | KTRI98 |
| 11-May-2004 | 36.97 | -86.56 | Warren | Kentucky | J. Schibig & M. Vance | KWAR1 |
| 11-May-2004 | 36.96 | -86.56 | Warren | Kentucky | J. Schibig & M. Vance | KWAR6 |
| 20-May-1964 | 34.22 | -89.36 | Lafayette | Mississippi | B. W. Haglan | MISS0018293 |
| 20-Oct-2000 | 34.37 | -89.29 | Lafayette | Mississippi | B. A. Connolly & M. Duffy | MISS0018290 |
| 18-Sep-1980 | 34.43 | -89.39 | Lafayette | Mississippi | K. L. Gordon | MMNS006950 |
| 3-Nov-1965 | 34.54 | -89.49 | Lafayette | Mississippi | L. Dowdy | MISS0018292 |
| 29-Jul-1955 | 34.24 | -88.62 | Lee | Mississippi | J. D. Ray, Jr. | MISSA014060 |
| 13-Sep-2005 | 33.44 | -88.32 | Lowndes | Mississippi | M. Whitson | KNK3197300000000 |
| 11-Jul-1969 | 34.17 | -88.00 | Marion | Alabama | R. Kral | KANU212195 |
| 29-Sep-1967 | 34.66 | -89.47 | Marshall | Mississippi | S. Lassetter | MISS0018297 |
| 26-Jun-2005 | 34.32 | -89.47 | Lafayette | Mississippi | M. Alexander | MLAF1 |
| 28-Jun-1955 | 33.89 | -88.25 | Monroe | Mississippi | J. D. Ray, Jr. | MISSA014059 |
| 13-May-1993 | 34.00 | -88.57 | Monroe | Mississippi | R. G. Wieland | MMNS026542 |
| 10-May-1952 | 34.05 | -88.56 | Monroe | Mississippi | R. M. Schuster | MISSA014058 |
| 26-Jun-1980 | 33.34 | -89.63 | Montgomery | Mississippi | K. L. Gordon | MMNS006740 |
| 1-Sep-1968 | 36.44 | -87.49 | Montgomery | Tennessee | J. Schibig | NCU00153931 |
| 6-Dec-2004 | 34.59 | -88.19 | Tishomingo | Mississippi | Maddox, C. Neel | MTIS1 |
| 4-Aug-1996 | 34.24 | -89.22 | Pontotoc | Mississippi | C. T. Bryson | MMNS012849 |
| 4-Aug-1996 | 34.24 | -89.19 | Pontotoc | Mississippi | C. T. Bryson | MISS0018298 |
| 24-Jun-1951 | 33.57 | -86.45 | Saint Clair | Alabama | C. G. Hollis | UNA00032086 |
| 20-Aug-1962 | 33.94 | -86.26 | Saint Clair | Alabama | P. E. Bostick | NCU00153908 |
| 13-Oct-2018 | 35.18 | -89.29 | Fayette | Tennessee | R.G. Laport, D. Smith & J. Ng | SWMT05626 |
| 13-Oct-2018 | 35.16 | -89.29 | Fayette | Tennessee | R.G. Laport, D. Smith & J. Ng | SWMT05627 |
| 18-Jul-1966 | 36.57 | -87.97 | Stewart | Tennessee | B. Evans | NCU00153932 |
| 9-May-1980 | 34.12 | -90.07 | Tallahatchie | Mississippi | R. A. Stewart II | MMNS004476 |
| 20-Jul-2002 | 35.45 | -86.27 | Bedford | Tennessee | C. Neel | TBED1 |
| 8-Oct-2005 | 35.43 | -86.25 | Bedford | Tennessee | C. Neel | TBED14 |
| 1-Jan-2004 | 35.36 | -86.53 | Bedford | Tennessee | C. Neel | TBED4 |
| 1-Dec-2003 | 35.87 | -85.98 | Cannon | Tennessee | C. Neel | TCAN1 |
| 5-Aug-2005 | 35.80 | -85.99 | Cannon | Tennessee | M. Vance | TCAN36 |
| 9-Jan-2005 | 35.89 | -88.46 | Carroll | Tennessee | J. Schibig & M. Vance | TCAR1 |
| 8-Oct-2005 | 35.49 | -86.14 | Coffee | Tennessee | C. Neel | TCOF3 |
| 18-Aug-2002 | 36.07 | -86.82 | Davidson | Tennessee | J. Schibig | TDAV4 |
| 14-Sep-2002 | 36.27 | -86.90 | Davidson | Tennessee | J. Schibig | TDAV7 |
| 15-Jul-2003 | 36.09 | -85.82 | Dekalb | Tennessee | J. Schibig & J. H. Craddock | TDEK15 |
| 4-Aug-2002 | 36.06 | -85.86 | Dekalb | Tennessee | J. Schibig, Barber, J. H. Craddock & Torkelson | TDEK4 |
| June-02 | 35.04 | -89.23 | Fayette | Tennessee | B. Turner | TFAY25 |
| 1-May-2002 | 35.04 | -89.24 | Fayette | Tennessee | B. Turner | TFAY6 |
| 20-Jul-2002 | 35.28 | -86.14 | Franklin | Tennessee | C. Neel | TFRA1 |
| 25-Sep-2005 | 35.07 | -86.05 | Franklin | Tennessee | P. Tarpley | TFRA3 |
| 25-Sep-2005 | 35.08 | -86.03 | Franklin | Tennessee | P. Tarpley | TFRA4 |
| 25-Sep-2005 | 35.07 | -86.02 | Franklin | Tennessee | P. Tarpley | TFRA5 |
| 25-Sep-2005 | 35.07 | -86.03 | Franklin | Tennessee | P. Tarpley | TFRA6 |
| 6-Jun-2000 | 35.38 | -87.19 | Giles | Tennessee | J. Schibig | TGIL1 |
| 2-Jun-1998 | 35.08 | -87.22 | Giles | Tennessee | C. Neel | TGIL3 |
| 1-Jun-1998 | 35.09 | -87.22 | Giles | Tennessee | C. Neel | TGIL4 |
| 24-Feb-2002 | 35.80 | -87.58 | Hickman | Tennessee | J. Schibig, Barber & J. H. Craddock | THIC1 |
| 2-Aug-2005 | 36.36 | -87.62 | Houston | Tennessee | J. Schibig & M. Vance | THOU1 |
| 1-Feb-2003 | 35.96 | -87.91 | Humphreys | Tennessee | J. Schibig & M. Vance | THUM1 |
| 28-Oct-2006 | 36.18 | -87.79 | Humphreys | Tennessee | J. Schibig, Leach, Greenwell & Dreaden | THUM4 |
| 15-Sep-1982 | 34.84 | -88.973 | Tippah | Mississippi | D. Meeks | MMNS013370 |
| 10-Sep-1978 | 34.53 | -88.16 | Tishomingo | Mississippi | K. E. Rogers | MMNS000789 |
| 15-Nov-2003 | 34.61 | -88.17 | Tishomingo | Mississippi | H. M. Sullivan | MMNS022035 |
| 3-Jun-2006 | 34.61 | -88.28 | Tishomingo | Mississippi | L. Majure & G. N. Ervin | MISSA014053 |
| unknown | 34.74 | -88.24 | Tishomingo | Mississippi | A. Allison | MISS0018300 |
| 5-May-2015 | 34.81 | -88.19 | Tishomingo | Mississippi | T. G. Harbison, M. T. Perkins, J. H. Craddock | NCU00085298 |
| 4-Oct-1978 | 34.89 | -88.11 | Tishomingo | Mississippi | K. E. Rogers | MMNS007370 |
| 5-Oct-1978 | 34.92 | -88.17 | Tishomingo | Mississippi | K. E. Rogers | MMNS002655 |
| 21-Jul-2002 | 35.31 | -86.54 | Lincoln | Tennessee | C. Neel | TLIN1 |
| 21-Jul-2002 | 35.04 | -86.56 | Lincoln | Tennessee | C. Neel | TLIN2 |
| 20-Jul-2002 | 35.03 | -86.47 | Lincoln | Tennessee | C. Neel | TLIN3 |
| 19-Jun-2004 | 36.59 | -85.86 | Macon | Tennessee | J. Schibig & M. Vance | TMAC1 |
| 8-Oct-2005 | 36.49 | -85.94 | Macon | Tennessee | J. Schibig, M. Vance & Torkelson | TMAC2 |
| 20-Jul-2002 | 34.67 | -86.52 | Madison | Tennessee | C. Neel | TMAD1 |
| 2-Oct-1999 | 34.66 | -86.51 | Madison | Tennessee | C. Neel | TMAD3 |
| 5-Jan-2002 | 35.34 | -86.78 | Marshall | Tennessee | J. Schibig, C. Neel & Torkelson | TMAR1 |
| 1-Oct-1998 | 35.46 | -87.12 | Maury | Tennessee | Estes | TMAU1 |
| 20-Jul-2002 | 35.40 | -86.30 | Moore | Tennessee | C. Neel | TMOO1 |
| 15-Oct-2004 | 35.54 | -87.98 | Perry | Tennessee | C. Neel | TPER1 |
| 7-Jan-2006 | 35.63 | -87.91 | Perry | Tennessee | C. Neel | TPER2 |
| 7-Jan-2006 | 35.68 | -87.88 | Perry | Tennessee | C. Neel | TPER3 |
| 8-Jul-2004 | 36.04 | -86.60 | Davidson | Tennessee | M. Vance | TPUT2 |
| 9-Jun-2002 | 36.44 | -86.74 | Robertson | Tennessee | Torkelson | TROB3 |
| 5-Jan-2003 | 36.44 | -86.75 | Robertson | Tennessee | Torkelson | TROB4 |
| 14-Mar-2003 | 35.71 | -86.22 | Rutherford | Tennessee | C. Neel | TRUT1 |
| 23-Jan-2003 | 36.40 | -85.88 | Smith | Tennessee | J. Schibig, Draper & Kemp | TSMI1 |
| 26-Jun-2003 | 36.21 | -85.87 | Smith | Tennessee | M. Vance, J. Schibig & J. H. Craddock | TSMI2 |
| 26-Jun-2002 | 36.62 | -88.03 | Stewart | Tennessee | J. Schibig | TSTE1 |
| 5-Sep-2002 | 36.57 | -88.01 | Stewart | Tennessee | J. Schibig | TSTE13 |
| 5-Sep-2002 | 36.57 | -88.02 | Stewart | Tennessee | J. Schibig | TSTE17 |
| 5-Aug-2002 | 36.61 | -88.04 | Stewart | Tennessee | J. Schibig | TSTE2 |
| 12-Sep-2002 | 36.61 | -88.04 | Stewart | Tennessee | J. Schibig | TSTE37 |
| 14-Jul-2003 | 36.66 | -88.05 | Stewart | Tennessee | J. H. Craddock | TSTE47 |
| 24-Jul-2003 | 36.67 | -88.05 | Stewart | Tennessee | J. H. Craddock | TSTE72 |
| 12-Sep-2003 | 36.60 | -88.01 | Stewart | Tennessee | J. H. Craddock | TSTE73 |
| 23-Apr-2002 | 36.51 | -86.45 | Sumner | Tennessee | J. Schibig | TSUM10 |
| 28-Apr-2002 | 36.51 | -86.39 | Sumner | Tennessee | J. Schibig | TSUM17 |
| 28-Apr-2002 | 36.43 | -86.68 | Sumner | Tennessee | J. Schibig | TSUM18 |
| 23-Oct-2003 | 36.38 | -86.66 | Sumner | Tennessee | J. Schibig | TSUM21 |
| 22-Jul-2004 | 36.62 | -86.21 | Sumner | Tennessee | J. Schibig & M. Vance | TSUM23 |
| 20-Nov-2003 | 36.30 | -88.66 | Weakley | Tennessee | J. Schibig & B. Turner | TWEA1 |
| 2-Feb-2003 | 35.83 | -87.00 | Williamson | Tennessee | J. Schibig | TWIL1 |
| 13-Jul-2004 | 35.82 | -87.06 | Williamson | Tennessee | J. Schibig & M. Vance | TWIL13 |
| 31-Aug-1996 | 33.20 | -89.04 | Winston | Mississippi | J. R. MacDonald | MMNS024435 |
| 31-Aug-1996 | 33.21 | -89.06 | Winston | Mississippi | J. R. MacDonald | MISSA014056 |
| 11-Oct-1978 | 33.21 | -88.97 | Winston | Mississippi | K. L. Gordon | MMNS005245 |
| 10-May-1967 | 34.02 | -87.36 | Winston | Alabama | R. C. Clark | NCU00153909 |
| 14-May-1999 | 33.87 | -89.57 | Yalobusha | Mississippi | R. A. Stewart II | MMNS034421 |
| 12-Jun-1996 | 33.98 | -89.82 | Yalobusha | Mississippi | J. R. MacDonald | MMNS012824 |
| 9-Apr-2020 | 34.40 | -86.20 | Marshall | Alabama | michaelezell | iNaturalist |
| 20-Apr-2020 | 34.74 | -86.71 | Madison | Alabama | N. Cash | iNaturalist |
| 29-May-2019 | 35.06 | -85.84 | Marion | Tennessee | N. Parrish | iNaturalist |
| 3-Aug-2019 | 35.06 | -88.73 | McNairy | Tennessee | A. J. Trently | iNaturalist |
| 9-Jul-2019 | 35.21 | -85.86 | Marion | Tennessee | dmp83178 | iNaturalist |
| 1-Jul-2020 | 35.56 | -87.45 | Lewis | Tennessee | M. Mayfield | iNaturalist |
| 17-May-2020 | 36.07 | -86.90 | Davidson | Tennessee | N. Parrish | iNaturalist |
| 16-Sep-2019 | 36.89 | -86.95 | Logan | Kentucky | rustyduck | iNaturalist |
| 14-Apr-2020 | 36.97 | -86.82 | Logan | Kentucky | rustyduck | iNaturalist |

#### Environmental layers

We downloaded bioclim environmental variables (1970-2000 averages) plus elevation at 30 arcsecond resolution (~1km^2^ at the equator) from Worldclim (ver. 2.1; www.worldclim.org; Hijmans *et al.*, 2005; Fick and Hijmans, 2017). We additionally downloaded the 2016 CONUS land cover and tree canopy cover datasets for the United States from the Multi-Resolution Land Characteristics Consortium (https://www.mrlc.gov; Yang *et al.*, 2018). All environmental variable layers were scaled to the same pixel resolution (~1km^2^; 0.00833° × 0.00833°) and cropped in QGIS (ver. 3.10.9; www.qgis.org) to span an area encompassing our occurrence records, which were used as “projection” layers for our models (above; Table 1; Fig. 1). We created additional independent sets of “calibration” environmental layers by creating mask layers comprising 5km, 10km, and 30km radius buffers surrounding each retained chestnut occurrence record in QGIS before trimming the original environmental layers with these buffer masks. The three sets of “calibration” layers were used to evaluate the set of environmental parameters that best predicted the observed occurrences (see Model Calibration below). We evaluated sets of environmental layers trimmed to 5km, 10km, and 30km buffers around each occurrence record in an effort to generate environmental niche models that appropriately accounted for the current “patchy” occurrences, very low reproductive output, and limited dispersal ability of remnant *C. dentata* when models were projected to the “projection” layers (Paillet, 2002; Dalgleish *et al.*, 2016). Moreover, the current deciduous forest habitat in the lower Mississippi River Valley and adjacent areas is fragmented, with high-quality upland forest patches that could harbor *C. dentata* often being widely dispersed and adjacent to unsuitable agricultural fields, silviculture, road corridors, and/or suburban development (Heilman *et al.*, 2002; Riitters *et al.*, 2012; Yang *et al.*, 2018).

**Figure 1.**
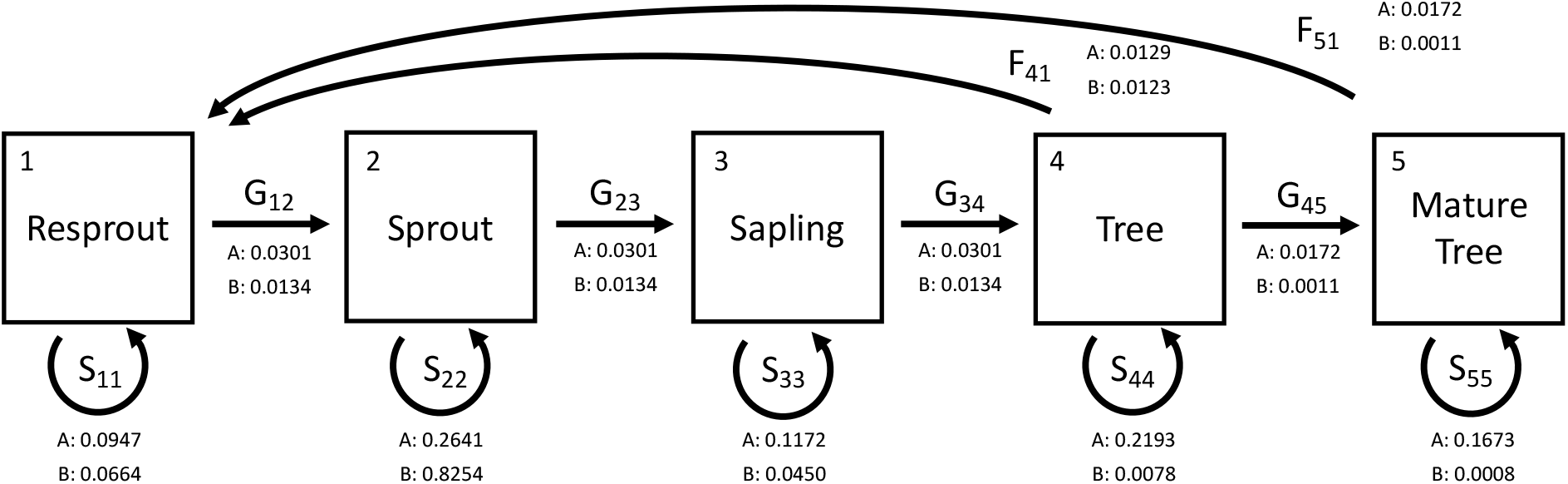
Life cycle diagram for a stage-classed *Castanea dentata* population in W Tennessee, N Mississippi, SW Kentucky, and NW Alabama. All trees were assigned to stage classes based on diameter at breast height (DBH) measurements. Stage classes are represented by squares (Resprout (stage 1) ≤ 1cm DBH, Sprout (stage 2) = 1.1-2.5cm DBH, Sapling (stage 3) = 2.6-10cm DBH, Tree (stage 4) = 10.1-20cm DBH, Mature Tree (stage 5) ≥ 20cm DBH). Arrows represent growth (G) transitions between stages, stasis (S) within a stage class, and fecundity (F) of reproductive stage classes. Values associated with transitions are elasticities for a model without blight (A) and a model with blight (B), representing the proportional importance of changes to a transition/stasis/fecundity rate on population growth

We performed correlation analyses on the calibration environmental layers using ENMTools (ver. 1.3; Warren *et al.*, 2010) to reduce overfitting of the distribution model to the occurrence points. Using an iterative approach, we removed environmental variables that exhibited Pearson correlation coefficients ≥ 0.7 with another variable before running the correlation analysis again to further evaluate pairwise correlations. When removing correlated variables, we retained one variable of each correlated pair/set during each iteration of the correlation analysis that is thought to be important for the life history of American chestnut based upon prior research (*e.g.*, Griscom and Griscom, 2012; Fei *et al.*, 2012; Santoro, 2013; Barnes and Delborne, 2019). Additionally, we excluded mean temperature of the wettest quarter (Bio 8), mean temperature of the driest quarter (Bio 9), precipitation of the warmest quarter (Bio 18), and precipitation of the coldest quarter (Bio 19) as prior analyses indicate these environmental layers may contain some apparent artifacts that can bias model construction (*e.g.*, Escobar *et al.*, 2014; Campbell, 2015). We retained a combination of environmental variables that exhibited low correlation coefficients (Pearson correlation coefficients ≤ 0.7) and/or are important life history determinants for American chestnut for model calibration: annual mean temperature (Bio 1), mean diurnal range (Bio 2), temperature seasonality (Bio 4), temperature annual range (Bio 7), annual precipitation (Bio 12), precipitation of wettest quarter (Bio 16), elevation, land cover, and canopy cover (Griscom and Griscom, 2012; Fei *et al.*, 2012; Santoro, 2013; Barnes and Delborne, 2019).

#### Model calibration

We conducted model calibration using the maximum entropy ecological niche modeling application Maxent (ver. 3.4.1; Philips *et al.*, 2006; Philips *et al.*, 2020), following the procedure of Simões *et al.* (2020) to identify the combination of model parameters and environmental variables that best fit our occurrence data. Maxent relies only on georeferenced presence data to create complex models using machine learning to relate environmental variables to occurrence points (Philips *et al.*, 2006; Merow *et al.*, 2013). We created 34,782 candidate models (11,594 candidate models each for the 5km, 10km, and 30km buffer sets) reflecting unique combinations of the retained environmental variables (see above), regularization multipliers (0.1, 0.3, 0.5, 0.7, 0.9, 1, 2, 3, 4, 5, 6), and feature classes (l, q, p, t, h, lq, lp, lt, lh, qp, qt, qh, pt, ph, th, lqp, lqt, lqh, lpt, lph, lth, qpt, qph, qth, pth, lqpt, lqph, lqth, lpth, qpth, lqpth, where l = linear, q = quadratic, p = product, h = hinge). The candidate models were implemented in Maxent using the R (ver. 4.0.2, R Core Team 2020) package kuenm (Cobos *et al.*, 2019) using an R script modified from the example file in Simões *et al.* (2020; Supplemental File 1). We randomly chose 80% of the 140 chestnut occurrence points for training the candidate models and 20% of the occurrence points for testing the candidate models. Candidate model performance was evaluated by partial ROC to evaluate the predictive fit of the model to the data via the receiver operating characteristic curve (ROC; Peterson *et al.*, 2008), omission rates (Anderson *et al.*, 2003), and model complexity (Akaike Information Criterion corrected for small sample size; AICc; Warren and Seifert, 2011).

Thirty candidate models met the statistical retention criteria of partial ROC, omission rate, and AICc (Simões *et al.*, 2020; Table 2). We implemented all of the models passing statistical criteria in Maxent (5km buffer = 22 models, 10km buffer = 7 models, 30km buffer =1 model) and projected them to the “projection” layers with clamping and Multivariate Environmental Similarity Surface (MESS) metrics. For all models, we evaluated variability in model predictions using 10 random bootstrap resampling replicates of 75% of the occurrence points (reserving 25% of occurrence points for testing each model replicate) in Maxent. To estimate land area with predicted environmental suitability for *C. dentata* we thresholded each of the median model outputs (cloglog format, WGS84 (EPSG:4326) projection) from Maxent to consider areas with an environmental suitability ≥0.5 as potential areas of occurrence (=1). Model areas with suitability <0.5 were considered environmentally unsuitable for occurrence (= 0).

**Table 2.**
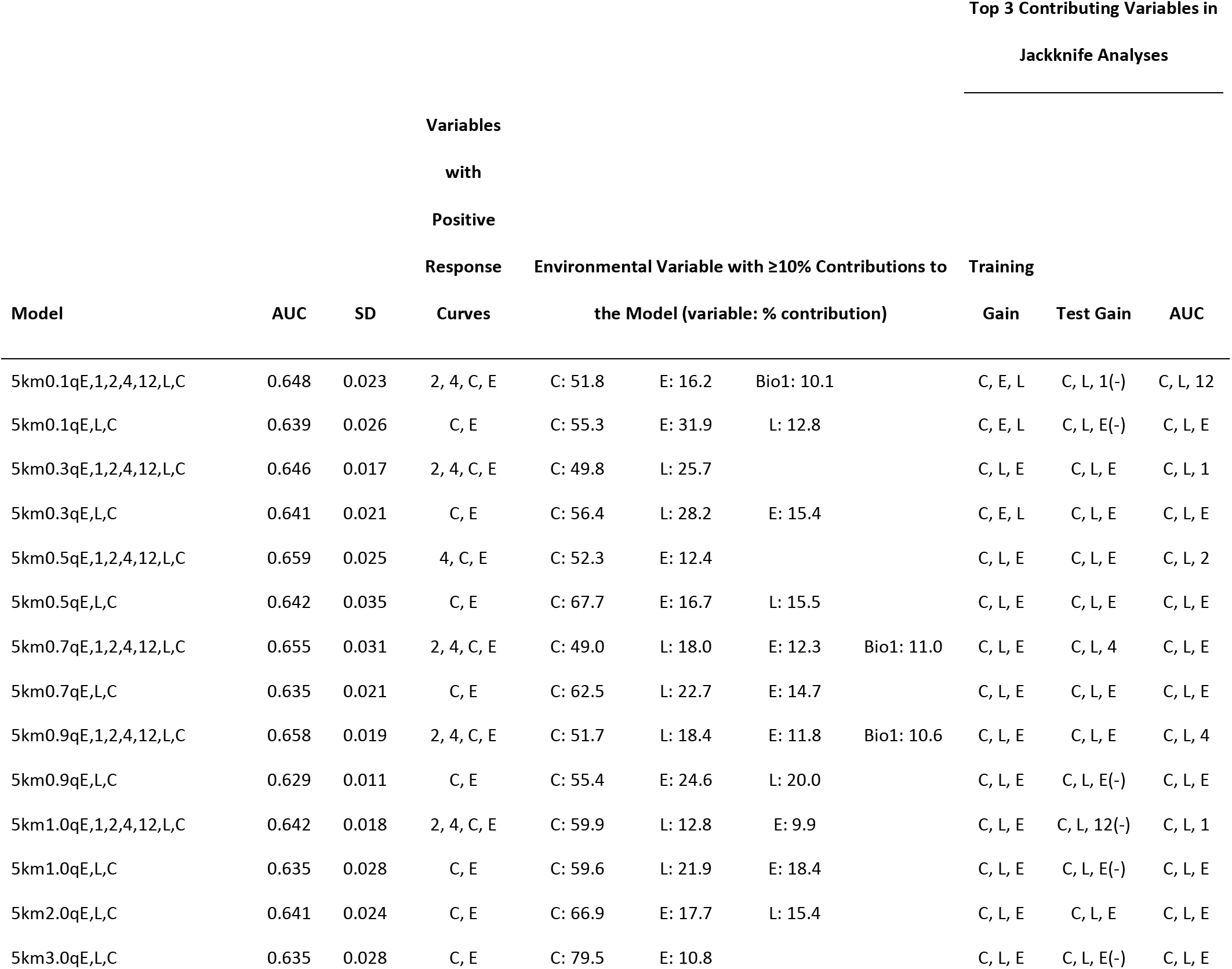
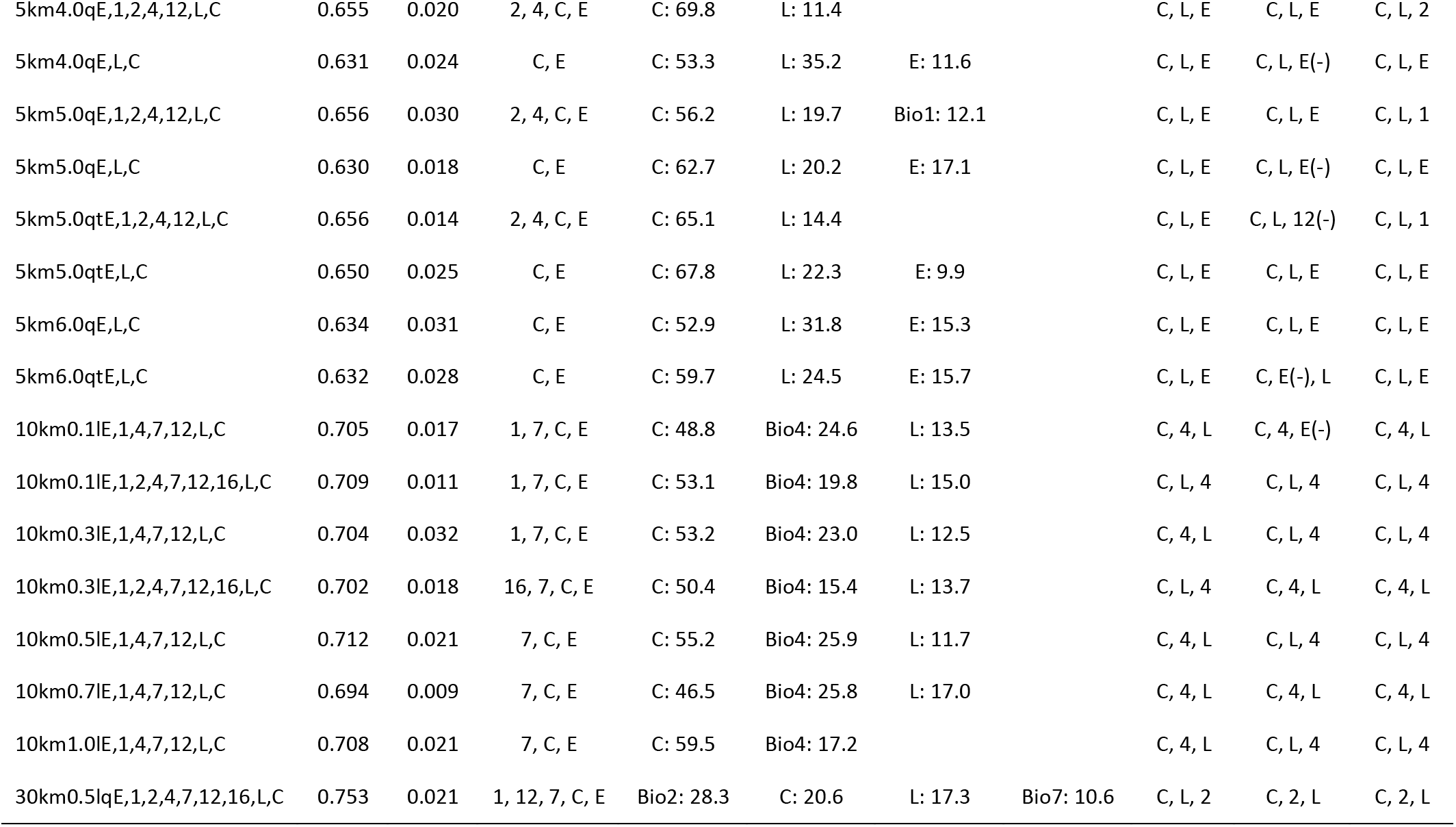
Summary statistics of the 30 environmental niche models passing model calibration criteria after implementation in Maxent. Model names take the form: buffer layer size (5km, 10km, or 30 km), regularization multiplier (ranging from 0.1-6.0), feature classes (l = linear, q = quadratic), and environmental variables (E = elevation, L = land cover, C = canopy cover, 1 = Bio 1, 2 = Bio 2, 4 = Bio 4, 7 = Bio 7, 12 = Bio 12, 16 = Bio 16). Environmental variable abbreviations are the same for all columns. Variables with negative model effects are indicated with a “-” in parentheses

### DEMOGRAPHIC MODEL

#### Tree occurrence data

To model the population demography of chestnut in W Tennessee, N Mississippi, SW Kentucky, and NW Alabama we filtered the dataset within the area of interest (NW corner: 37.3583°, −91.1444°, SE corner: 32.8416°, −85.6044°) to include any observations that included information on tree size, demography, and/or blight infection status. For example, when explicit height or stem diameter measurements (*e.g.*, Diameter at Breast Height, DBH) were not available in the primary source data, we retained occurrences that included notations such as “resprout,” “sapling,” or “small tree.” In total our thinned dataset comprised 558 occurrences that include information on tree size (Supplemental File 2), and we assigned each occurrence to one of five size classes representing life history transitions: resprouts (0-1.0 cm DBH), sprouts (1.1-2.5 cm DBH), saplings (2.5-10.0 cm DBH), trees (10.1-25.0 cm DBH), and mature trees (>25.0 cm DBH). Because of their prevalence in the dataset, trees noted as having a stem diameter “<1 cm” were assigned a DBH of 0.5 cm and retained as resprouts. This resulted in a “current” population structure of 23.7% resprouts, 47.5% sprouts, 19.7% saplings, 6.8% trees, and 2.3% mature trees. For modeling, we assumed that all of these trees were genetically independent, rather than treating spatially clustered occurrences as arising from a single root crown, and assumed that historical records were part of a “current” population throughout the area of interest. Blight affliction status among tree size classes was treated as a binary state (no blight = 0, blight = 1) based upon first-hand observation or collector notations. From these observations/notations we computed the proportion of trees in each size class that were blight-free and applied these proportions to the entire dataset. Blight-affliction was assumed to be lethal and result in tree mortality without size class regressions.

We modeled population growth rate, stable size class distributions, and the importance of different life history stages on American chestnut population growth rates in the area of interest using a stage structured Lefkovitch (1965) projection matrix model (Caswell, 2001; Morris and Doak, 2002). Our approach multiplies a transition matrix containing estimated probabilities of stasis/transition between demographic stages with a vector describing the observed distribution of individuals in the five demographic stages (Table 3a). To model the effect of blight on the chestnut population, we multiplied the transition rates in our transition matrix by the proportion of blight-free trees in each size class, thereby proportionally reducing the transition rates in the transition matrix to produce a “blight” matrix (Table 3b). We modeled the demography of the population (Fig. 1) under two different scenarios: 1) without blight, providing an expectation for the demographic future of the species in the absence of blight in the area of interest, and 2) with the inclusion of blight to examine the demographic future of the species in the area of interest under current conditions. To parameterize our transition matrix (Table 3), we modified individual transition rates between size classes from two prior studies of American chestnut demography with trees in similar size classes to our study: one near the historical northeastern range limit in Maine where chestnut was not strongly affected by blight (Elwood, 2014), and one focused on naturalized populations in Michigan exhibiting a range of blight affliction (Davelos and Jarosz, 2004). Where size regression was parameterized in prior studies, we assumed stage class stasis because we lacked life history information from our observations to confidently include these rates. Both models were implemented in R (ver. 4.0.0) using the package ‘popbio’ (Stubben and Milligan, 2007) for 100 time steps. Prior demographic studies of American chestnut have estimated each time step to represent multiple years (*e.g.*, ~13 years; Elwood, 2014; Van Drunen *et al.*, 2017). To summarize the output of our models we obtained the dominant eigenvalues representing λ (where λ = 1 represents a stable population size, λ < 1 represents population contraction, and λ > 1 represents population growth) and the projected stable size class distribution. We also estimated sensitivity and elasticity values from each model to infer the importance of different life history stages and transition rates on population growth rate.

**Table 3.** Transition matrices showing estimated stasis (on diagonal), growth (below diagonal), and reproductive (upper right) elements A) without blight and B) with blight for a population of American chestnut in W Tennessee, N Mississippi, SW Kentucky, and NW Alabama. Matrix elements were derived from rates estimated for populations of American chestnut in S Maine (Elwood, 2014) and Michigan (Davelos and Jarosz, 2004). Size class regressions in prior studies were included in rates for size class stasis

| **A. w/o Blight** | **1. Resprout** | **2. Sprout** | **3. Sapling** | **4. Tree** | **5. Mature Tree** |
| --- | --- | --- | --- | --- | --- |
| **1. Resprout** | 0.826 | 0 | 0 | 0.693 | 2.167 |
| **2. Sprout** | 0.171 | 0.977 | 0 | 0 | 0 |
| **3. Sapling** | 0 | 0.023 | 0.866 | 0 | 0 |
| **4. Tree** | 0 | 0 | 0.134 | 0.957 | 0 |
| **5. Mature Tree** | 0 | 0 | 0 | 0.043 | 0.987 |
| **B. w/ Blight** | **1. Resprout** | **2. Sprout** | **3. Sapling** | **4. Tree** | **5. Mature Tree** |
| **1. Resprout** | 0.765 | 0 | 0 | 0.693 | 2.167 |
| **2. Sprout** | 0.159 | 0.905 | 0 | 0 | 0 |
| **3. Sapling** | 0 | 0.021 | 0.709 | 0 | 0 |
| **4. Tree** | 0 | 0 | 0.110 | 0.338 | 0 |
| **5. Mature Tree** | 0 | 0 | 0 | 0.015 | 0.380 |

## Results

### ENVIRONMENTAL NICHE MODELS

The 5km buffer “calibration” layers (*i.e.*, 5km buffer around each occurrence point) produced ENMs having AUCs (area under the curve of the receiver operating characteristic) ranging from 0.629 (SD +/− 0.011) to 0.659 (SD +/− 0.025). Model response curves for Bio 2, Bio 4, canopy cover, and elevation showed positive relationships with the predicted probability of presence in models including variables in addition to elevation, canopy cover, and landcover. In models including only elevation, canopy cover, and land cover as explanatory variables, canopy cover and elevation showed positive relationships with the predicted probability of presence. Canopy cover (49.0%-79.5%), elevation (9.9%-31.9%), and land cover (11.4%-31.8%) made the largest contributions to most of the models with Bio 1 (10.1%-12.1%) contributing to some models. In Maxent jackknife analyses, canopy cover, land cover, and elevation were the most important for training gain, test gain, and AUC with Bio 1, Bio 2, Bio 4, and Bio 12 being important for test gain and AUC in a few models (Table 2).

With the 10km buffer “calibration” layers (*i.e.*, 10km buffer around each occurrence point) the resulting ENMs had AUCs ranging from 0.694 (SD +/− 0.009) to 0.712 (SD +/− 0.021). Model response curves for Bio 7, canopy cover, and elevation showed positive relationships with the predicted probability of presence in most models, with Bio 1 and Bio 16 having positive relationships in some models. Canopy cover (46.5-59.5%), Bio 4 (15.4-25.9%), and land cover (11.7-15.0%) made the largest contributions to the models. In Maxent jackknife analyses, canopy cover, land cover, and Bio 4 were the most important for training gain, test gain, and AUC, with elevation being important for test gain in one model (Table 2).

The 30km buffer “calibration” layer (*i.e.*, 30km buffer around each occurrence point) produced an ENM with an AUC of 0.753 (SD +/− 0.021). Model response curves for Bio 1, Bio 7, Bio 12, canopy cover, and elevation all showed positive relationships with the predicted probability of presence. Bio 2 (28.3%), canopy cover (20.6%), and land cover (17.3%) made the largest contributions to the model, and in Maxent jackknife analyses, canopy cover, land cover, and Bio 2 were most important for training gain, test gain, and AUC (Table 2).

Projecting the ENMs to the “projection” layers indicated relatively high environmental suitability for American chestnut throughout much of W Tennessee, N Mississippi, SW Kentucky, and NW Alabama (Fig. 2). The 30 model predictions were roughly visually similar, though many models differed appreciably in areas of predicted environmental suitability (Supplemental File 3). The areas of highest environmental suitability were most consistently predicted to occur throughout eastern portions of the area of interest (*e.g.*, Hardeman, McNairy, Chester, Henderson, Hardin, Decatur, and Benton Counties in Tennessee; Yalobusha, Lafayette, Marshall, Benton, Tippah, Union, Alcorn, Prentiss, Tishomingo, Itawamba, and Monroe Counties in Mississippi; Lauderdale, Colbert, Franklin, Marion, and Lamar Counties in Alabama) that comprise higher elevations and areas of high canopy cover (as inferred from the elevation and canopy cover layers downloaded from WorldClim and MRLCC, respectively). However, areas of relatively high predicted environmental suitability also occurred along river drainages and uplands otherwise surrounded by low elevation areas of agriculture in the western portion of the area of interest between the Tennessee R. and the Mississippi R. (Fig. 2).

**Figure 2.**
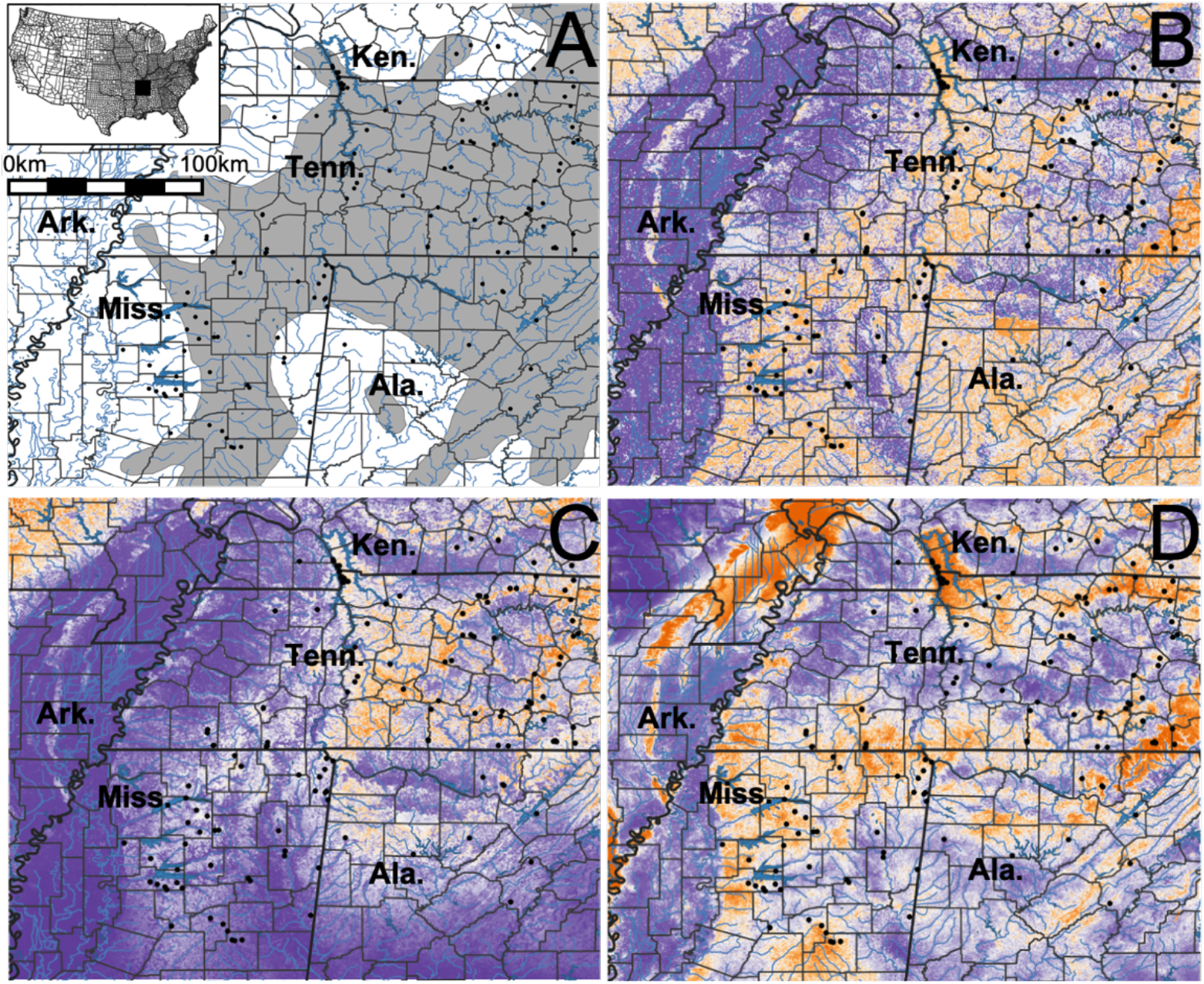
A) The estimated historical range of *Castanea dentata* (gray shading; Little, 1977) in W Tennessee (Tenn.), N Mississippi (Miss.), SW Kentucky (Ken.), and NW Alabama (Ala.). The black square in the inset map indicates the study location. Representative ENM projections using B) 5km, C) 10km, and D) 30km environmental layer buffers around each point for model construction. In B, C, and D, warmer (*i.e.*, more orange) colors indicate higher predicted environmental suitability. The model projections resulting from the 5km buffers were generally more apparently “realistic” compared to those arising from the 10km and 30km buffers (Supplemental File 2). Black dots in all panels show the location of occurrence records within the area of interest that were used for ENM construction. County boundaries are indicated by the lighter borders within each state

Visual inspection of the top model outputs constructed from each of the environmental layer buffer sets suggested several of the models resulting from the 5km and 10km buffers produced the most realistic predictions for the potential distribution of *C. dentata* in the area of interest, as they apparently better reflected areas of high canopy cover (Fig. 1; Supplemental File 3). Models resulting from the 30km buffer (and several resulting from the 5km and 10km buffers) made either apparent under- or over-predictions of habitat suitability given the patchiness of available forest habitat throughout W Tennessee, N Mississippi, SW Kentucky, and NW Alabama (Supplemental File 3). Models that included other environmental variables in addition to elevation, canopy cover, and land cover tended to have broader predictions of environmental suitability than those relying only on elevation, canopy cover, and land cover. Models resulting from the 10km buffer tended to underpredict suitability in N Mississippi relative to the models resulting from the 5km buffer. All of the models also predicted areas of high environmental suitability in parts of E Arkansas comprising uplands and areas of high canopy cover (*e.g.*, Crowley’s Ridge, foothills of E Ozark Mountains) beyond the western extreme of the accepted distribution. After thresholding the models resulting from the 5km buffers, approximately 14,589,787ha (range: 5,529,005-28,464,967ha) of land area, representing ~47% of the total projection area, was predicted to contain environmentally suitable habitat (environmental suitability ≥0.5) for *C. dentata*. From the 10km buffers approximately 3,532,389ha (range: 21,187-9,700,077ha) of land area, representing ~11% of the total projection area, was predicted to represent environmentally suitable habitat (environmental suitability ≥0.5). Approximately 9,856,276ha of land area, representing ~32% of the total projection area, was predicted as environmentally suitable for American chestnut by the model resulting from the 30km buffer.

### DEMOGRAPHIC MODEL

Without blight, the model for the American chestnut population in W Tennessee, N Mississippi, SW Kentucky, NW Alabama produced a dominant eigenvalue (λ) of 1.09 representing a growing population. Projecting the population over 100 time steps produced exponentially growing trajectories for all size classes (Fig. 3). The projected stable size class distribution indicated a population comprising 34.2% resprouts, 52.5% sprouts, 5.4% saplings, 5.6% trees, 2.4% mature trees. Sensitivities were largest for the transition between sprouts and saplings, and for the transition between trees and mature trees indicating life-history transitions from small stages to mid-size stages, and to reproductive maturity, are important for population growth in the absence of blight. Summed elasticities were higher for matrix elements representing stasis (S = 0.863) than for elements representing growth (G = 0.107) or fecundity (F = 0.030; Fig. 2) indicating stage class stasis is also important for population growth in the absence of blight.

**Figure 3.**
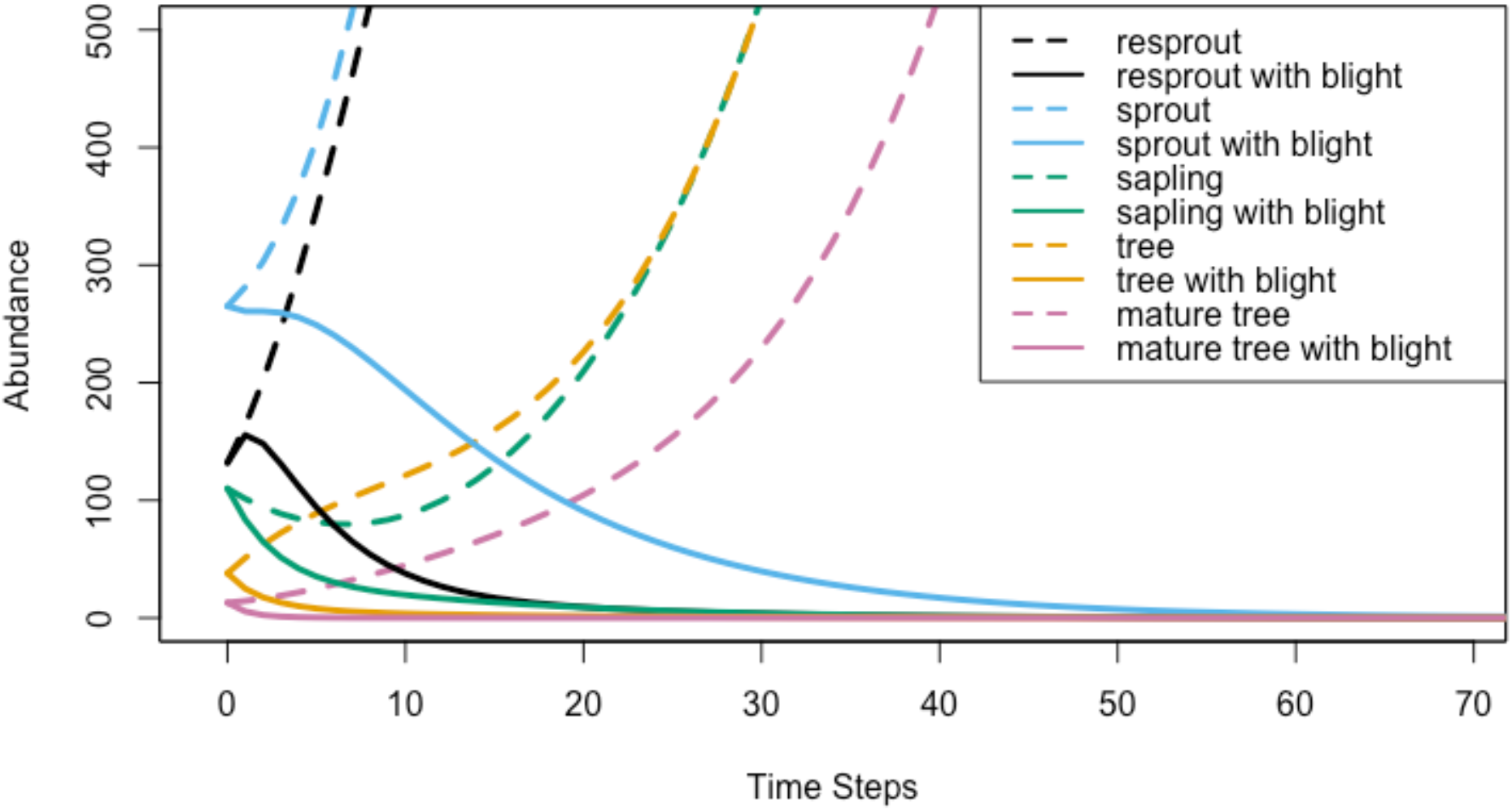
Demographic model projections for a *Castanea dentata* population in W Tennessee, N Mississippi, SW Kentucky, and NW Alabama without blight (dashed lines) and with blight (solid lines). Each line represents the demographic trajectory for a different size class as indicated in the inset key. Without blight, the population was predicted to have a λ = 1.09 indicating an increase ~9% per time step, while with blight the population is predicted to have a λ = 0.92 indicating a decrease of ~8% per time step. The duration of each time step is not known for the study population, but is estimated to extend over multiple years for a similar population in southern Ontario, Canada (~13 years; Van Drunen *et al.*, 2018), suggesting extinction of American chestnut throughout the study region could occur in ~100-200 years

After proportionally reducing the transition rates between stage classes by accounting for only blight-free trees, the model produced a dominant eigenvalue (λ) of 0.92 indicating a shrinking population. Projecting the population over 100 time steps produced rapidly declining trajectories for all size classes (Fig. 3). The projected stable size class distribution indicated a population comprising 7.7% resprouts, 82.4% sprouts, 8.3% saplings, 1.6% trees, and 0.04% mature trees. Sensitivities were largest for the stasis of sprouts, and for the transition between sprouts and saplings (Fig. 2) indicating stasis and transition from small stages to mid-size stages is important for population growth in the presence of blight. Summed elasticities were higher for matrix elements representing stasis (S = 0.945) than for elements representing growth (G = 0.041) or fecundity (F = 0.013) also indicating that in the presence of blight persistence is important for population growth.

## Discussion

The functional extinction of *C. dentata* after the introduction of *C. parasitica* in the early 1900s is thought to have drastically altered the ecology and community dynamics of eastern North American forests (Jacobs *et al.*, 2013, Dalgleish *et al.*, 2016). Unfortunately, the rapid near-extirpation of an integral component of eastern deciduous forest communities before many formal ecological theories and methods were established, makes current efforts to reconstruct pre-blight *C. dentata* ecology, and the consequences of its loss from eastern forests, a difficult task. For example, historical records give some indication of pre-blight forest composition (Zon, 1904; Mattoon, 1909; Ashe, 1912; Brooks, 1937; Russell, 1987; Paillet, 2002) and how these forests may have responded to the loss of *C. dentata* (Ziegler, 1920; Keever, 1953; Stephenson *et al.*, 1991; Jacobs *et al.*, 2013), but it remains unclear whether such descriptions can be extrapolated throughout the large and ecologically variable historical range of *C. dentata*. Studies after blight spread throughout the historical range of American chestnut have inferred significant community-level consequences, including replacement of chestnut by other tree species (*e.g.*, *Quercus* spp., *Carya* spp. and *Acer rubrum*; Keever, 1953; Stephenson *et al.*, 1991; Davelos and Jarosz, 2004; Griscom and Griscom, 2012), and the probable extinction of interacting species (*e.g.*, greater chestnut weevil; *Curculio caryatrypes* Boheman 1843; Anderson, 2017). Yet, *C. dentata* persists throughout much of its historical range as a minor component of eastern forests comprising small individuals that continue to re-sprout from the root systems of blight-killed trees, which typically succumb to blight prior to reaching reproductive size, though occasional reproductive trees have also been noted (*e.g.*, Paillet, 1984; Boucher, 2000; Dalgleish *et al.*, 2016). Recent efforts to characterize the extent of remaining suitable habitat likely to harbor extant *C. dentata* have made it clear that additional effort is needed to locate remnant individuals/populations and to more fully characterize the current ecological and demographic status of the species throughout its formerly native range to aid conservation and restoration efforts (Fei *et al.*, 2007; Santoro, 2013; Jacobs *et al.*, 2013; Dalgleish *et al.*, 2016). In this study, we focused on an area of the historical American chestnut range that has been relatively understudied to identify potential new areas in which remnant populations of *C. dentata* may occur but remain undiscovered, and to gain insight into the current and future population dynamics of remnant trees in an area with high genetic diversity that could bolster breeding and restoration efforts by increasing the heterozygosity of breeding stock (Stilwell *et al.*, 2003).

### ENVIRONMENTAL NICHE MODELS

Prior applications of environmental niche modeling to *C. dentata* have sought to investigate the ecology of American chestnut and identify suitable habitat for discovering remnant populations, as well as to explore areas suitable for reintroduction. Some of these studies have focused on restricted portions of the historical range, carefully investigating the environmental parameters associated with the presence of *C. dentata* in areas of relatively intact forest (Fei *et al.*, 2007; Santoro, 2013; Elwood, 2014). Predictions from these studies indicate that suitable habitat for *C. dentata* in Shenandoah National Park occurs in primarily well-drained upland forests at relatively high elevations in the Blue Ridge Mountains of Virginia (Santoro, 2013) and in areas of high canopy cover on steep upper- and mid-slope areas overlying sandstone in topographically diverse parts of Mammoth Cave National Park in Kentucky with young forests on previously cultivated lands being unsuitable for chestnut occurrence (Fei *et al.*, 2007). At a continental scale that sacrifices microscale variation and nuance, environmental suitability is predicted to shift northward and beyond the current northern range limit in response to climate change (Fei *et al.*, 2012; Barnes and Delborne, 2019). However, none of these studies have included many occurrences from the southwestern distribution of American chestnut and offer only limited insight into environmental suitability near the historical southwestern range limit.

Though our study is focused on a relatively restricted portion of American chestnut’s historical range we found that much of W Tennessee, N Mississippi, SW Kentucky, and NW Alabama remain environmentally suitable and that many potentially undiscovered *C. dentata* may persist in remaining forests of the region. The 30 environmental niche models satisfying statistical test criteria produced modest (AUC <0.9), roughly similar, but widely variable habitat suitability predictions throughout the area of interest (Fig. 1; Supplemental File 2). Over-predictions seemed apparent when models included bioclimatic variables compared to those that only included elevation, canopy cover, and land cover, while under-predictions seemed apparent with some sets of environmental layer buffers (Supplemental File 2). The variability among model predictions likely reflects, in part, the difficulty in producing a multivariate model that captures the environmental niche represented by the disparate environments where chestnut observations occur (Vale *et al.*, 2014; Barnes and Delborne, 2019). Prior to the introduction of chestnut blight, *C. dentata* was one of the most broadly distributed tree species in eastern North America, ranging from southern Maine to southern Mississippi (Peattie, 1950; Little, 1977; Paillet, 2002; Jacobs *et al.*, 2013; Dalgleish *et al.*, 2016). Given this historical distribution, the species likely also occupied a much broader set of ecological conditions than currently realized by extant populations (Fei *et al.*, 2007; Griscom and Griscom, 2012). Indeed, historical documents note the occurrence of the species at or beyond the currently accepted distributional limits in SW Tennessee (Fraser, 1864; Williams, 1873), though likely as relatively minor forest components (Ashe 1912). Although our environmental niche models incorporate historical observations of *C. dentata* from the time *C. parasitica* had arrived in W Tennessee and N Mississippi, well within the lifespan of healthy *C. dentata* (Zon, 1904; Ashe, 1912), they likely underestimate the full range of environmental conditions that could be occupied by the species. This may reflect non-representative historical reporting or collection of specimens due to sampling bias against a common species (prior to blight), but remnant populations may also occur in habitats where blight was slower to arrive. Moreover, populations at distributional limits are rare and may occupy environmental conditions not typically experienced by core distribution populations or be found in small patches of suitable environmental conditions embedded within unsuitable environments (Vale *et al.*, 2014; Angert, 2020). These possibilities would make it difficult to accurately identify the set of environmental parameters that reliably predict species occurrences (Galante *et al.*, 2018). Thus, most of our model predictions, especially those produced from the 5km buffers, likely represent conservative projections of suitable habitat (Fig. 1). Search efforts for remnant populations of *C. dentata* in under-explored parts of the historical distribution may prove fruitful, and intact forest with high predicted suitability beyond the currently accepted range should be considered as possibly viable habitat. In Mississippi, for example, apparently relatively little contemporary search effort has occurred, even though this area may harbor crucial genetic diversity for breeding and restoration efforts by representing potential genetic relicts that more closely reflect populations that expanded from refugia following the last glacial maximum (Russell, 1987; Kubisiak and Roberds, 2006; Dane, 2009). Interestingly, most of our models also predict suitable American chestnut habitat in upland areas of NE Arkansas. While observations of *C. dentata* are not known from these areas, some of these areas are habitat for the closely-related congeners *Castanea ozarkensis* Ashe and *Castanea pumila* Mill, but could potentially harbor rare *C. dentata*.

As with all environmental niche models, caution is warranted in interpreting these projections. Our model calibration approach produced several “equally-good” models that when implemented and projected to the area of interest produced widely differing predictions of environmental suitability. One challenge to producing accurate models of habitat suitability appears to be determining the scale at which environmental heterogeneity influences the presence of *C. dentata* (Fei *et al.*, 2007; Santoro, 2013). Because of adult mortality caused by chestnut blight, the population dynamics of American chestnut are likely not currently strongly influenced by pollen and seed dispersal, or seedling recruitment, and the probability of occurrence in many areas (including the southwestern historical range limit) is probably highest within or adjacent to intact forests that have persisted since the arrival of blight (*e.g.,* >50 years) with relatively low disturbance (Paillet, 1988; 2002; Fei *et al.*, 2007; Griscom and Griscom, 2012; Gustafson *et al.*, 2017). The frequency and intensity of *C. parasitica* affliction also plays a role in the occurrence and persistence of chestnut, and environmental variation over relatively small scales might strongly influence blight affliction (Van Drunen *et al.*, 2018). Thus, model construction using relatively small environmental layer “buffers” around each occurrence point may more accurately capture the environmental covariates of *C. dentata* occurrence and persistence over the last several decades.

Variability in niche model prediction might also be expected near a species range limit if the observed distribution represents the realized rather than the fundamental ecological niche (Warren, 2012; Broennimann *et al.*, 2021). The naturalization of *C. dentata* beyond the accepted historical range limit in Michigan, Wisconsin, Iowa, and Nova Scotia (Paillet and Rutter, 1989; Barnes and Delborne, 2019), suggests that *C. dentata* may not have been at ecological equilibrium when standardized range maps were established (Little, 1977) and may still have been expanding from glacial refugia before the arrival of chestnut blight (Russell, 1987; Huang *et al.*, 1998). Some of this expansion would likely have been contingent upon ongoing evolutionary responses (*i.e.*, mutation, fitness differences, selection, local adaptation, etc.) to changing and novel climatic conditions encountered during expansion, and may not be adequately accounted for in environmental niche modeling approaches that make predictions assuming niche conservatism (Weins and Graham, 2005). The accuracy of our environmental niche models may also have been affected by lowered habitat residency of *C. dentata* as a consequence of a root rot pathogen (*Phytophthora cinnamomi*) that is thought to have diminished the range extent and population densities of *C. dentata* in the southern US by the late 1800s (Russell, 1987; Anagnostakis, 2001; Dalgleish *et al.*, 2016). Environmental niche models relying upon post-*P. cinnamomi* and post-*C. parasitica* occurrences of American chestnut are likely less representative of the habitat once occupied by *C. dentata*, though further investigation is warranted to better understand how models might be impacted by pathogenic mortality.

### DEMOGRAPHIC MODEL

Demographic modeling of remnant chestnut populations can reveal important population dynamics that may guide conservation priorities and highlight the challenges of restoration efforts (Boucher, 2000; Davelos and Jarosz, 2004; Jacobs, 2007, 2013; Elwood, 2014; Baines *et al.*, 2014). With our simple models, we aimed to investigate the general population trajectories for remnant trees in W Tennessee, N Mississippi, SW Kentucky, and NW Alabama, a relatively understudied portion of the historical American chestnut range. After making several simplifying assumptions about a relatively modest sampling of individuals spread throughout the region across several decades we found that the small starting population was projected to either expand exponentially (λ >1) in the absence of blight or decline rapidly (λ <1) when blight is accounted for over the next several decades (Fig. 3). In the absence of blight we projected exponential growth of ~9% per time step across all size classes, while accounting for blight resulted in a projection of a ~8% decrease per time step across all size classes (Fig. 3).

In the modeled blight and non-blight scenarios, the high stasis element elasticities (Fig. 2) indicate individual tree persistence over many years as having the greatest proportional influence on population growth rates and that this trait has likely structured the current stage class distribution of remnant American chestnut in W Tennessee, N Mississippi, SW Kentucky, and NW Alabama. Approximately 70% of individuals in our observed data are represented by the smallest size classes (resprouts or sprouts). In the presence of recurrent *C. parasitica* infection these stage classes indicate regrowth from previously blight-afflicted individuals and are crucial for longer-term persistence of the species in relatively undisturbed habitat where blight-afflicted root crowns remain viable (Zon, 1904; Paillet, 1988). Large sensitivities for transitions to larger and mature trees in the demographic model also suggest that even rare trees transitioning to reproductive maturity can play an outsized role in population growth. In the model accounting for blight, the chestnut population in W Tennessee, N Mississippi, SW Kentucky, and NW Alabama is projected to decline rapidly toward extinction within ~20 time steps (~100-200 years) at minimum. However, the population could persist much longer if time steps are assumed to span a decade or longer (Van Drunen *et al.*, 2017). Yet, these individuals cannot persist indefinitely and will eventually experience mortality from blight, senescence, or stochastic environmental influences (*e.g.*, storms, tree fall, land development, etc.) without interventions to preserve them.

The actual population of extant trees in the area of interest is likely much larger than the observed set of trees in our dataset, and the actual stage transition rates for individuals in W Tennessee, N Mississippi, SW Kentucky, and NW Alabama likely differ from those in southern Maine and Michigan. Nevertheless, the model projections presented here are qualitatively similar to the demographic projections from other studies of *C. dentata* populations suggesting that currently the major determinant of chestnut population dynamics is *C. parasitica* and the presence of relatively undisturbed forest (Paillet, 1988; Elwood, 2014; Van Drunen *et al.*, 2017; 2018). Pre-blight estimates suggest *C. dentata* comprised up to 5-10% of forest trees west of the Tennessee R. (Ashe, 1912). Though current densities are certainly much lower than historical occurrences, the species was an important component of forest communities between the Mississippi and Tennessee Rivers, and continues to contribute to the ecological dynamics of forests in this area. However, we emphasize that our model projections should be considered with caution and serve as a starting point for more robust investigations of a species that has long attracted attention from land managers, biologists, and the general public (Kummer, 2003; Steiner and Carlson, 2006; Rosen, 2013; Popkin, 2020). Continued surveys to discover remnant trees and more consistent documentation of blight affliction, especially in under-explored areas, would help refine model projections while serving the dual purpose of aiding conservation efforts.

## Conclusions

Despite being nearly extirpated in the mid-20^th^ century by introduced chestnut blight, *C. dentata* continues to persist throughout its range primarily as small resprouting individuals. Recent research emphasis has been centered on core parts of the historical distribution where remnant trees appear to be plentiful. Much less effort has focused on the southwestern part of the range—where the species exemplifies its classification as a species of “Special Concern” (Tennessee Flora Committee, 2015; Tennessee Natural Heritage Program, 2016)—and the degree to which suitable habitat and undiscovered populations persist in under-surveyed parts of the historical distribution is not well known. We found that suitable habitat for *C. dentata* occurs throughout much of the historical southwestern range extent in W Tennessee, N Mississippi, SW Kentucky, and NW Alabama using environmental niche modeling parameterized from locations where American chestnut has been observed over the last several decades. However, variability among modest model predictions also suggests that the extreme western portions of the historical native range represent marginal habitat for the species and areas of highest environmental suitability are predicted to be restricted to higher elevations and current areas of high forest canopy cover (Fig. 1). Although W Tennessee, N Mississippi, SW Kentucky, and NW Alabama is a region that has been impacted over the last ~150 years by timber harvesting, forest fragmentation (Heilman *et al.*, 2002; Riitters *et al.*, 2012), and other introduced plant pathogens (*i.e.*, *Phytophthora cinnamomi*; Anagnostakis, 2001) our ENMs suggest the area has not supported very large populations of *C. dentata* in recent times. Demographic modeling using historical and contemporary observations of *C. dentata* across the region also supports this inference and suggests a dire near-term future for the persistence of the species near its southwestern range margin. The density of recently-documented American chestnut was generally low throughout the area of interest (but higher in some large areas of relatively intact forest), and demographic modeling accounting for the observed effects of blight suggests drastic declines across all size classes over the next ~100-200 years. Continued population decline appears to be the rule in the presence of *C. parasitica* given the devastating effects the fungus has had on *C. dentata* populations over the past ~120 years (Ziegler, 1920; Paillet, 2002; Davelos and Jarosz, 2004; Jacobs, 2007, 2013; Dalgleish *et al.*, 2016), though local climatic and habitat variability might moderate the effects of blight in some areas leaving relatively healthy populations (Van Drunen *et al.*, 2018). Taken together, our ENM and stage-structured demographic projections highlight the continued presence of suitable *C. dentata* habitat throughout the southwestern extent of its historical distribution, and emphasize the need for efforts to locate, conserve, and introduce genetic material from individuals with potentially novel and locally adapted genotypes from these areas into currently active restoration programs.

## Acknowledgements

The authors are grateful to D. Smith (Fayette County Forestry Association) and C. Neel (American Chestnut Foundation-Tennessee Chapter) for assistance with field collections and comments on a draft of this manuscript. The prior research of J. Schibig (Volunteer State Community College) on *C. dentata* in middle and west Tennessee was crucial in guiding our research efforts. S. Hasty and J. Porter (Rhodes College) provided crucial research support.

---

Supplemental File 1. R scripts used for environmental niche model calibration (adapted from Simões *et al.*, 2020) and demographic matrix modeling in ‘popbio’ (Stubben and Milligan, 2007).

Supplemental File 2. Locality and size information for all W Tennessee, N Mississippi, SW Kentucky, and NW Alabama American chestnut observations comprising the demographically modeled population. Observations included historical collections from herbaria, personal observations by the authors and colleagues, and citizen naturalist observations from GBIF recorded and verified with the iNaturalist application.

Supplemental File 3. Environmental niche model projections from Maxent across W Tennessee, N Mississippi, SW Kentucky, and NW Alabama for all 30 models passing model calibration criteria. The models arising from the 5km buffer calibration layers (the first 12 models) generally produced more apparently realistic projections for suitable habitat compared to the models arising from the 10km buffer calibration layers (next 7 models) or the 30km buffer calibration layers (last model). Many of the 10km buffer models underestimated parts of the N Mississippi range, while the 30km buffer model predicts suitable habitat in many areas where chestnut is not observed. Some of the 5km buffer models that incorporate Bioclim variables with elevation, land cover, and canopy cover also make apparent overpredictions of environmental suitability.

## Literature Cited

Anagnostakis, S. L. 2001. The effects of multiple importations of pests and pathogens on a native tree. Biol. Invasions, 3:245–254.

Angert, A. L., H. D. Bradshaw, Jr., and D. W. Schemske. 2008. Using experimental evolution to investigate geographic range limits in monkeyflowers. Evolution, 62:2660–2675.

Angert, A. L., M. G. Bontrager, and J. Ågren. 2020. What do we really know about adaptation at range edges? Annu. Rev. Ecol. Evol. S., 51:341–361.

Ashe, W. W. 1912. Chestnut in Tennessee. Tennessee Geological Survey Bulletin 10-B. Nashville, TN, U.S.A.

Anderson, R. 2017. Co-extinction and the case of American chestnut and the greater Chestnut weevil (*Curculio caryatrypes*). Canadian Museum of Nature Blog. https://canadianmuseumofnature.wordpress.com/2017/02/02/co-extinction-and-the-case-of-american-chestnut-and-the-greater-chestnut-weevil-curculio-caryatrypes/.

Anderson, R. P., D. Lew, and A. T. Peterson. 2003. Evaluating predictive models of species’ distributions: criteria for selecting optimal models. Ecol. Model., 162:211–232.

Baines, A. D., E. A. Eager, and A. M. Jarosz. 2014. Modeling and analysis of American chestnut populations subject to various stages of infection. Lett. Biomath., 1:235–247.

Barnes, J. C., and J. A. Delborne. 2019. Rethinking restoration targets for American chestnut using species distribution modeling. Biodivers. Conserv., 28:3199–3220.

Boucher, D. H. 2000. Population dynamics of American chestnut sprouts: the projection matrix approach. J. Am. Chestnut Found., 14:38–45.

Brooks, A. B. 1937. Castanea dentata. Castanea, 2:61–67.

Broennimann, O., B. Petitpierre, M. Chevalier, M. González-Suárez, J. M. Jeschke, J. Rolland, S. M. Gray, S. Bacher, and A. Guisan. 2021. Distance to native climatic niche margins explains establishment success of alien mammals. Nat. Commun., 12:2353 dx.doi.org//https://doi.org/10.1038/s41467-021-22693-0.

Campbell, L. P., C. Luther, D. Moo-Llanes, J. M. Ramsey, R. Danis-Lozano, and A. T. Peterson. 2015. Climate change influences on global distributions of dengue and chikungunya virus vectors. Philos. T. Roy. Soc. B, 370:1–9.

Caswell, H. 2001. Matrix Population Models: Construction, Analysis, and Interpretation, second edition. Sinauer, Sunderland, MA, U.S.A.

Cobos, M. E., A. T. Peterson, N. Barve, and L. Osorio-Olvera. 2019. kuenm: an R package for detailed development of ecological niche models using Maxent. PeerJ 7:e6281.

Dalgleish, H. J., and R. K. Swihart. 2012. American chestnut past and future: implications of restoration for resource pulses and consumer populations of eastern U.S. forests. Restor. Ecol., 20:490–497.

Dalgleish, H. J., C. D. Nelson, J. Scrivani, and D. Jacobs. 2016. Consequences of shifts in abundance and distribution of American chestnut for restoration of a foundation forest tree. Forests, 7 dx.doi.org//10.3390/f7010004.

Dane, F. 2009. Comparative phylogeography of *Castanea* species. Acta Hortic., 844:211–222.

Davelos, A. L., and A. M. Jarosz. 2004. Demography of American chestnut populations: effects of a pathogen and a hyperparasite. J. Ecol., 92:675–685.

De Bruijn, A., E. J. Gustafson, D. M. Kashian, H. J. Dalgleish, B. R. Sturtevant, and D. F. Jacobs. 2014. Decomposition rates of American chestnut (*Castanea dentata*) wood and implications for coarse woody debris pools. Can. J. Forest Res., 44:1575–1585.

Diamond, S. J., R. H. Giles, R. L. Kirkpatrick, and G. J. Griffin. 2000. Hard mast production before and after the chestnut blight. South. J. Appl. For., 24:196–201.

Elliot, K. J., and W. T. Swank. 2008. Long-term changes in forest composition and diversity following early logging (1919-1923) and the decline of American chestnut (*Castanea dentata*). Plant Ecol., 197:155–172.

Ellison, A. M., M. S. Bank, B. D. Clinton, E. A. Colburn, K. Elliott, C. R. Ford, D. R. Foster, B. D. Kloeppel, J. D. Knoepp, G. M. Lovett, J. Mohan, D. A. Orwig, N. L. Rodenhouse, W. V. Sobczak, K. A. Stinson, J. K. Stone, C. M. Swan, J. Thompson, B. Von Holle, and J. R. Webster. 2005. Loss of foundation species: consequences for the structure and dynamics of forested ecosystems. Front. Ecol. Environ., 3:479–486.

Ellstrand, N. C., and D. R. Elam. 1993. Population genetic consequences of small population size: implications for plant conservation. Annu. Rev. Ecol. Syst., 24:217–242.

Elwood, E. C. 2014. Population matrix model for American chestnut (*Castanea dentata*) and the implications for re-introduction [Honors thesis]. The College of William and Mary. Williamsburg, VA, U.S.A.

Escobar, L. E., A. Lira-Noriega, G. Medina-Vogel, and A. T. Peterson. 2014. Potential for spread of the white-nose fungus (*Pseudogymnoascus destructans*) in the Americas: use of Maxent and NicheA to assure strict model transference. Geospatial Health, 9:221–229.

Fei, S., J. Schibig, and M. Vance. 2007. Spatial habitat modeling of American chestnut at Mammoth Cave National Park. Forest Ecol. Manag., 252:201–207.

Fei, S., L. Liang, F. L. Paillet, K. C. Steiner, J. Fang, Z. Shen, Z. Wang, and F. V. Hebard. 2012. Modelling chestnut biogeography for American chestnut restoration. Divers. Distrib., 18:754–768.

Fick, S. E., and R. J. Hijmans. 2017. WorldClim 2: new 1km spatial resolution climate surfaces for global land areas. Int. J. Climatol., 37:4302–4315.

Fraser, H. 1864. Hattie Fraser Diary 1861-1864. Craddock Book Club, Somerville, TN, U.S.A.

Galante, P. J., B. Alade, R. Muscarella, S. A. Jansa, S. M. Goodman, and R. P. Anderson. 2018. The challenge of modeling niches and distributions for data-poor species: a comprehensive approach to model complexity. Ecography, 41:726–736.

Griscom, H. P., and B. W. Griscom. 2012. Evaluating the ecological niche of American chestnut for optimal hybrid seedling reintroduction sites in the Appalachian ridge and valley province. New Forest., 43:441–455.

Gustafson, E. J., A. De Bruijn, N. Lichti, D. F. Jacobs, B. R. Sturtevant, J. Foster, B. R. Miranda, and H. J. Dalgleish. 2017. The implications of American chestnut reintroduction on landscape dynamics and carbon storage. Ecosphere, 8:e01773.

Halbritter, A. H., R. Billeter, P. J. Edwards, and J. M. Alexander. 2015. Local adaptation at range edges: comparing elevation and latitudinal gradients. J. Evolution. Biol., 28:1849–1860.

Heilman Jr., G. E., J. R. Strittholt, N. C. Slosser, and D. A. Dellasala. 2002. Forest fragmentation of the conterminous United States: Assessing forest intactness through road density and spatial characteristics. BioScience, 52:411–422.

Hijmans, R. J., S. E. Cameron, J. L. Parra, P. G. Jones, and A. Jarvis. 2005. Very high resolution interpolated climate surfaces for global land areas. Int. J. Climatol., 25:1965–1978.

Huang, H. W., F. Dane, and T. L. Kubisiak. 1998. Allozyme and RAPD analysis of the genetic diversity and geographic variation in wild populations of the American chestnut (Fagaceae). Am. J. Bot., 85:1013–1021.

Jacobs, D. F. 2007. Toward development of silvical strategies for forest restoration of American chestnut (*Castanea dentata*) using blight-resistant hybrids. Biol. Conserv., 137:497–506.

Jacobs, D. F., H. J. Dalgleish, and C. D. Nelson. 2013. A conceptual framework for restoration of threatened plants: the effective model of American chestnut (*Castanea dentata*) reintroduction. New Phytol., 197:378–393.

Keever, C. 1953. Present composition of some stands of the former oak-chestnut forest in the southern Blue Ridge Mountains. Ecology, 34:44–55.

Kubisiak, T. L., and J. Roberds. 2006. Genetic structure of American chestnut populations based on neutral DNA markers. p. 109–122. *In*: Steiner, K.C. and J.E. Carlson (eds.). Restoration of American chestnut to forest lands—Proceedings of a conference and workshop. May 4-6, 2004, The North Carolina Arboretum. Natural Resources Report NPS/NCR/CUE/NRR-2006/001. National Park Service, Washington, D.C., U.S.A.

Kummer, C. 2003. A new chestnut. The Atlantic, 1 June 2003. https://www.theatlantic.com/magazine/archive/2003/06/a-new-chestnut/302742/?utm_source=copy-link&utm_medium=social&utm_campaign=share.

Laport, R. G. 2020. Remnant American chestnut (*Castanea dentata* (Marsh.) Borkh.; Fagaceae) in upland forests of western New York. Proc. Rochester Acad. Sci., 21:5–14.

Laport, R. G., D. Smith, and J. Ng. 2020. Remnant American chestnut (*Castanea dentata*) near the historical western range limit in southwestern Tennessee. Castanea, 85:232–243.

Lefkovitch, L. P. 1965. The study of population growth in organisms grouped by stages. Biometrics, 21:1–18.

Little, E. L., Jr. 1977. Minor eastern hardwoods. Volume 4. Atlas of United States trees. Miscellaneous Publication 1342. USDA Forest Service, Washington, D.C., U.S.A.

Mattoon, W. R. 1909. The origin and early development of chestnut sprouts. Forest Quarterly, 7:34–37.

Mccormick, J. F., and R. B. Platt. 1980. Recovery of an Appalachian forest following the chestnut blight. Am. Midl. Nat., 104:264–273.

Merow, C., M. J. Smith, and J. A. Silander, Jr. 2013. A practical guide to MaxEnt for modeling species’ distributions: what it does, and why inputs and settings matter. Ecography, 36: 1058–1069.

Morris, W. F., and D. F. Doak. 2002. Quantitative Conservation Biology: Theory and Practice of Population Viability Analysis. Sinauer, Sunderland, MA, U.S.A.

Paillet, F. L. 1982. The ecological significance of American chestnut (*Castanea dentata* (Marsh.) Borkh.) in the Holocene forests of Connecticut. B. Torrey Bot. Club, 109:457–473.

Paillet, F. L. 1988. Character and distribution of American chestnut sprouts in southern New England woodlands. B. Torrey Bot. Club, 115:32–44.

Paillet, F. L. 2002. Chestnut: history and ecology of a transformed species. J. Biogeogr., 29:1517–1530.

Paillet, F. L., and P. A. Rutter. 1989. Replacement of native oak and hickory tree species by the introduced American chestnut (*Castanea dentata*) in southwestern Wisconsin. Can. J. Botany, 67:3457–3469.

Peattie, D. 1950. A Natural History of Trees of Eastern and Central North America. Houghton Mifflin Company, Boston, MA, U.S.A.

Perrier, A., D. Sánchez-Castro, and Y. Willi. 2020. Expressed mutational load increases toward the edge of a species’ geographic range. Evolution, 74:1711–1723.

Peterson, A. T., M. Papeş, and J. Soberón. 2008. Rethinking receiver operating characteristic analysis applications in ecological niche modeling. Ecol. Model., 213:63–72.

Phillips, S. J., R. P. Anderson, and R. E. Schapire. 2006. Maximum entropy modeling of species geographic distributions. Ecol. Model., 190:231–259.

Phillips, S. J., M. Dudík, and R. E. Schapire. 2020. Maxent software for modeling species niches and distributions (Version 3.4.1). URL http://biodiversityinformatics.amnh.org/open_source/maxent/, accessed 15 August 2020.

Popkin, G. 2020. Can genetic engineering bring back the American chestnut? The New York Times Magazine, 30 April 2020. https://www.nytimes.com/2020/04/30/magazine/american-chestnut.html?smid=url-share.

R CORE TEAM. 2020. R: A language and environment for statistical computing. R Foundation for Statistical Computing, Vienna, Austria. URL https://www.R-project.org/

Riitters, K. H., and J. D. Wickham. 2012. Decline of forest interior conditions in the conterminous United States. Sci. Rep., 2:653 dx.doi.org//10.1038/srep00653.

Rosen, R. 2013. Genetically engineering and icon: can biotech bring the chestnut back to America’s forests? The Atlantic, 31 May 2013. https://www.theatlantic.com/technology/archive/2013/05/genetically-engineering-an-icon-can-biotech-bring-the-chestnut-back-to-americas-forests/276356/?utm_source=copy-link&utm_medium=social&utm_campaign=share.

Russell, E. W. B. 1987. Pre-blight distribution of *Castanea dentata* (Marsh.) Borkh. Bull. Torrey Bot. Club, 114:183–190.

Santoro, J. 2013. American chestnut (*Castanea dentata*) habitat modeling: identifying suitable sites for restoration in Shenandoah National Park, Virginia [M.Sc. thesis]. Duke University. Durham, NC, U.S.A.

Savolainen, O., T. Pyhäjärvi, and T. Knürr. 2007. Gene flow and local adaptation in trees. Annu. Rev. Ecol. Evol. S., 38:595–619.

Schibig, J., C. Neel, M. Hill, M. Vance, and J. Torkelson. 2005. Ecology of American chestnut in Kentucky and Tennessee. J. Am. Chestnut Found., 19:42–48.

Schibig, J., Vance, S. Cuming, L. Fly, C. Neel, and J. Torkelson. 2006. Ecology of the American chestnut and Allegheny chinquapin on the Cumberland Plateau of Kentucky and Tennessee. J. Am. Chestnut Found., 20:44–50.

Schwadron, P. A. 1995. Distribution and persistence of American chestnut sprouts, *Castanea dentata* (Marsh.) Borkh., in northeastern Ohio woodlands. Ohio J. Sci., 95:281–288.

Shaw, J., J. H. Craddock, and M. A. Binkley. 2012. Phylogeny and phylogeography of North American *Castanea* Mill. (Fagaceae) using cpDNA suggests gene sharing in the southern Appalachians. Castanea, 77:186–211.

Simões, M., D. Romero-Alvarez, C. Nuñez-Penichet, L. Jiménez, and M. E. Cobos. 2020. General Theory and Good Practices in Ecological Niche Modeling: A Basic Guide. Biodiversity Informatics, 15:67–68.

Smith, D. M. 2000. American chestnut: Ill-fated monarch of the eastern hardwood forest. J. Forest., 98:12–15.

Steiner, K. C. and Carlson, J. E, eds. 2006. Restoration of American chestnut to forest lands: proceedings of a conference and workshop. May 4-6, 2004, The North Carolina Arboretum. Natural Resources Report NPS/NCR/CUE/NRR 2006/001. National Park Service, Washington, D.C., U.S.A.

Stephens, G. R. and P. E. Waggoner. 1980. A half century of natural transitions in mixed hardwood forests. Bulletin 783. The Connecticut Agricultural Experiment Station, New Haven, CT, U.S.A.

Stephenson, S. L., H. S. Adams, and M. L. Lipford. 1991. The present distribution of chestnut in the upland forest communities of Virginia. Bull. Torrey Bot. Club, 118:24–32.

Stevens, D. L., K. Soltau, A. Davelos Baines, and A. M. Jarosz. 2014. American chestnut sprout dynamics. Acta Hortic., 1019:223–227.

Stilwell, K. L., H. M. Wilbur, C. R. Werth, and D. R. Taylor. 2003. Heterozygote advantage in the American chestnut, *Castanea dentata* (Fagaceae). Am. J. Bot., 90:207–213.

Stubben, C., and B. Milligan. 2007. Estimating and analyzing demographic models using the popbio package in R. J. Stat. Softw., 22:1–23.

TENNESSEE FLORA COMMITTEE. 2015. Guide to the Vascular Plants of Tennessee. University of Tennessee Press, Knoxville, TN, U.S.A.

TENNESSEE NATURAL HERITAGE PROGRAM. 2016. Rare Plant List. Department of Environment and Conservation, Nashville, TN, U.S.A.

Tindall, J. R., J. A. Gerrath, M. Melzer, K. Mckendry, B. C. Husband, and G. J. Boland. 2004. Ecological status of American chestnut (*Castanea dentata*) in its native range in Canada. Can. J. Forest Res., 34:2554–2563.

Vale, C. G., P. Tarroso, and J. C. Brito. 2014. Predicting species distribution at range margins: testing the effects of study area extent, resolution and threshold selection in the Sahara– Sahel transition zone. Divers. Distrib., 20:20–33.

Van Drunen, S. G., K. Schutten, C. Bowen, G. J. Bolen, and B. C. Husband. 2017. Population dynamics and the influence of blight on American chestnut at its northern range limit: Lessons for conservation. Forest Ecol. Manag., 400:375–383.

Van Drunen, S. G., J. L. Mccune, and B. C. Husband. 2018. Distribution and environmental correlates of fungal infection and host tree health in the endangered American chestnut in Canada. Forest Ecol. Manag., 427:60–69.

Warren, D.L. 2012. In defense of ‘niche modeling.’ Trends Ecol. Evol., 27:497–500.

Warren, D.L., and S. N. Seifert. 2011. Ecological niche modeling in Maxent: The importance of model complexity and the performance of model selection criteria. Ecol. Appl., 21:335–342.

Warren, D.L., R. E. Glor, and M. Turelli. 2010. ENMTools: a toolbox for comparative studies of environmental niche models. Ecography, 33:607–611.

Wiens, J. J, and C. H. Graham. 2005. Niche conservatism: Integrating evolution, ecology, and conservation biology. Annu. Rev. Ecol. Evol. S., 36:519–539.

Williams, J. S. 1873. Old times in west Tennessee: reminiscences, semi-historic, of pioneer life and the early emigrant settlers in the Big Hatchie country. W. G. Cheeney, Memphis, TN, U.S.A.

Yang, L., S. Jin, P. Danielson, C. Homer, L. Gass, S. M. Bender, A. Case, C. Costello, J. Dewitz, J. Fry, M. Funk, B. Granneman, G. C. Liknes, M. Rigge, and G. Xian. 2018. A new generation of the United States National Land Cover Database: Requirements, research priorities, design, and implementation strategies. ISPRS J. Photogramm., 146:108–123.

Ziegler, E.A. 1920. Problems arising from the loss of our chestnut. Forest Leaves, 17:152–155.

Zon, R. 1904. Chestnut in southern Maryland. U.S. Department of Agriculture, Bureau of Forestry, Bulletin No. 53. Washington, D.C., U.S.A.

