## Supplemental File 3 for "Environmental niche and demographic modeling of American chestnut near its southwestern range limit"

Supplemental File 3. Environmental niche model projections from Maxent across W Tennessee, N Mississippi, SW Kentucky, and NW Alabama for all 30 models passing model calibration criteria. The models arising from the 5km buffer calibration layers (the first 12 models) generally produced more apparently realistic projections for suitable habitat compared to the models arising from the 10km buffer calibration layers (next 7 models) or the 30km buffer calibration layers (last model). Many of the 10km buffer models underestimated parts of the N Mississippi range, while the 30km buffer model predicts suitable habitat in many areas where chestnut is not observed. Some of the 5km buffer models that incorporate Bioclim variables with elevation, land cover, and canopy cover also make apparent overpredictions of environmental suitability.

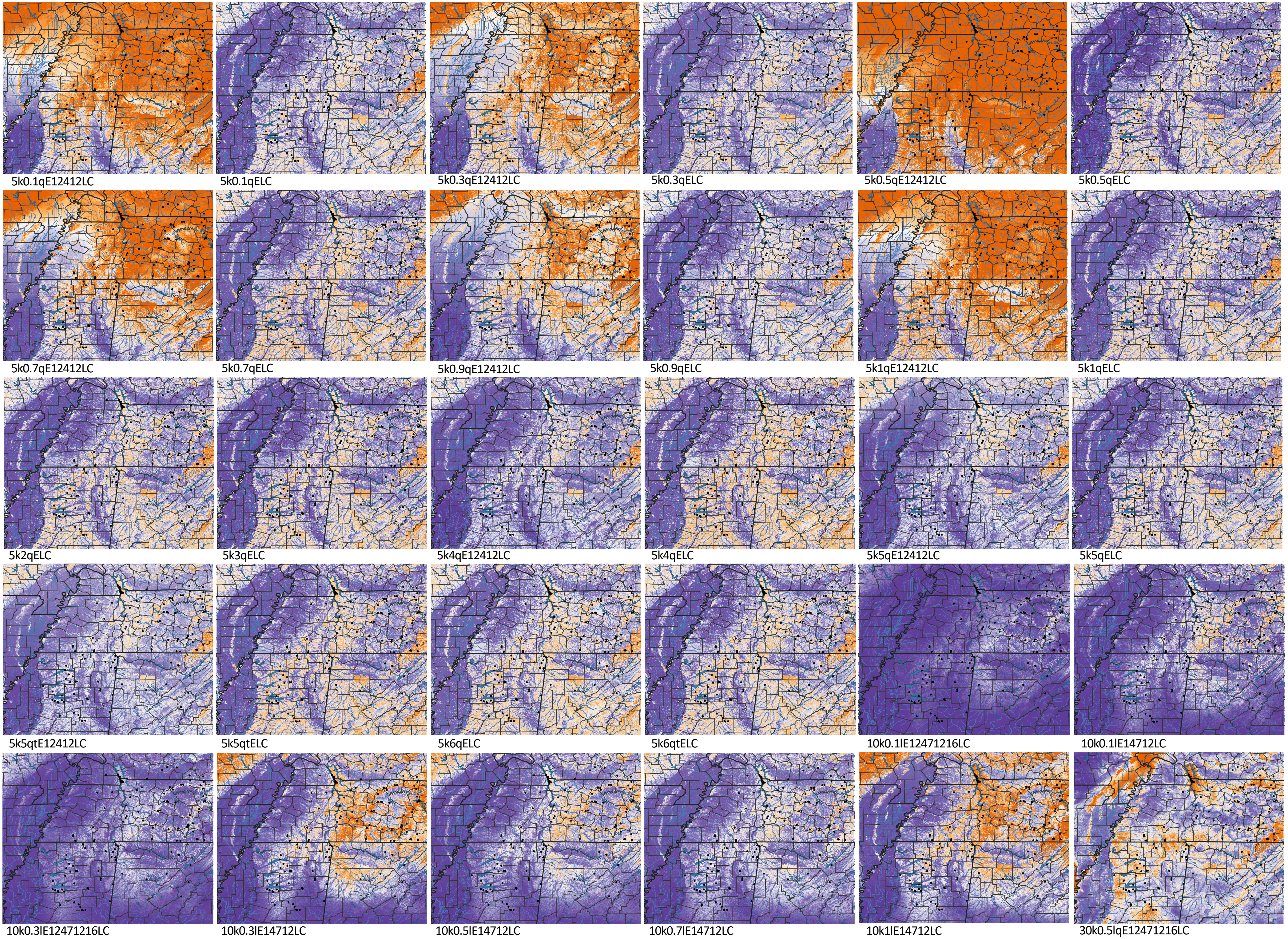
